# Quantifying changes in individual-specific template-based representations of center-of-mass dynamics during walking with ankle exoskeletons using Hybrid-SINDy

**DOI:** 10.1101/2022.01.31.478514

**Authors:** Michael C. Rosenberg, Joshua L. Proctor, Katherine M. Steele

**Affiliations:** Department of Mechanical Engineering, University of Washington, Seattle, United States of America; Department of Applied Mathematics, University of Washington, Seattle, United States of America

## Abstract

Ankle exoskeletons alter whole-body walking mechanics, energetics, and stability by altering center-of-mass (CoM) motion. Controlling the dynamics governing CoM motion is, therefore, critical for maintaining efficient and stable gait. However, how CoM dynamics change with ankle exoskeletons is unknown, and how to optimally model individual-specific CoM dynamics, especially in individuals with neurological injuries, remains a challenge. Here, we evaluated individual-specific changes in CoM dynamics in unimpaired adults and one individual with post-stroke hemiparesis while walking in shoes-only and with zero-stiffness and high-stiffness passive ankle exoskeletons. To identify optimal sets of physically interpretable mechanisms describing CoM dynamics, termed *template signatures*, we leveraged hybrid sparse identification of nonlinear dynamics (Hybrid-SINDy), an equation-free data-driven method for inferring sparse hybrid dynamics from a library of candidate functional forms. In unimpaired adults, Hybrid-SINDy automatically identified spring-loaded inverted pendulum-like template signatures, which did not change with exoskeletons (p>0.16), except for small changes in leg resting length (p<0.001). Conversely, post-stroke paretic-leg rotary stiffness mechanisms increased by 37-50% with zero-stiffness exoskeletons. While unimpaired CoM dynamics appear robust to passive ankle exoskeletons, how neurological injuries alter exoskeleton impacts on CoM dynamics merits further investigation. Our findings support Hybrid-SINDy’s potential to discover mechanisms describing individual-specific CoM dynamics with assistive devices.

## Introduction

Ankle exoskeletons are used to improve walking function and gait mechanics^1-3^. Personalized passive and powered ankle exoskeletons have shown promise to improve walking function and muscle coordination in some unimpaired adults and individuals with neurological injuries^2,4,5^. However, changes in gait in response to ankle exoskeletons are highly individualized, especially following neurological injury, making device personalization critical to improving mobility. For example, in stroke survivors and children with cerebral palsy, ankle exoskeletons elicit diverse—and sometimes detrimental—impacts on gait mechanics, walking speed, step length, and the energetic cost of walking^1-4^. Quantifying and characterizing these responses remain challenging and hinders clinicians’ and designers’ abilities to customize exoskeletons to support walking function.

Characterizing the changes in the neural and biomechanical processes governing center-of-mass (CoM) motion (*i.e*., CoM dynamics) with ankle exoskeletons may help explain heterogeneous exoskeleton impacts on task-level goals during walking. Despite observed changes in center-of-mass (CoM) mechanics and energetics with ankle exoskeletons, little is known about how CoM dynamics change with ankle exoskeletons to achieve task-level goals, like walking stably and efficiently^2,5-10^. CoM energetics are altered in unimpaired adults walking with ankle exoskeletons and following neurological injuries (*e.g*., post-stroke, the paretic leg exhibits reduced power generation and changes in leg power with exoskeletons differ between individuals)^2,5,11^. However, whether these changes in leg power are accompanied by changes in CoM dynamics or simply altered CoM kinematics is unclear.

Reduced-order representations of CoM dynamics, often termed *template* models, provide a foundation to quantify complex exoskeleton responses using interpretable mechanical elements^12-17^. For example, Full & Kodistchek (1999) used template models of CoM dynamics, such as the spring-loaded inverted pendulum (SLIP), to quantify strategies to stabilize the CoM in response to perturbations^15^. Such reduced-order representations of gait encode neural and biomechanical dynamics using a minimalist set of physics-based mechanisms. Specifically, common template models of CoM dynamics use a variety of mechanisms, such as SLIP leg springs or the rigid legs of an inverted pendulum walker, to describe relationships between leg kinematics and CoM accelerations during gait^10,13,15,16,18-22^. Each mechanism within a model, therefore, encodes a hypothesis about how neural and biomechanical subsystems interact to achieve task-level walking goals.

However, determining which template mechanisms are needed to optimally describe an individual’s CoM dynamics remains challenging. Inverted pendulum templates have been useful in modeling CoM energetics, the transition from walking to running^16,18^, and strategies for energetically efficient CoM acceleration^8,22,23^ and lateral stabilization^10,24^. Higher-dimensional template structures, such as the bipedal SLIP, were needed to more accurately predict sagittal-plane ground reaction forces (GRFs)^20,25^. Additional mechanisms applied to the bipedal SLIP template, such as leg dampers, rotary springs, or curved feet, have been used to further improve the accuracy of CoM dynamics models during walking^19,26^. This breadth of templates proposed for human walking suggests that the mechanisms that best describe gait are individual-specific^20,26^. Selecting individual-specific template structures (*i.e*., the mechanisms included in the template) is, therefore, critical to quantifying exoskeleton impacts on CoM dynamics using template models.

To emphasize the individual-specific nature of templates, we denote the combination of mechanisms best describing individual-specific CoM dynamics as a *template signature*. Inter- or intra-individual differences in the template signature mechanisms that best describe CoM dynamics for an individual and walking condition, or the coefficients estimated for each mechanism, may provide insight into how exoskeletons impact CoM dynamics. For example, template signatures in children with hemiparetic cerebral palsy differed from typically developing children and were asymmetric, with increased stiffness—defined by the coefficient of the stiffness mechanism—in the paretic leg^27,28^. Atypical and asymmetric CoM dynamics suggested that, following neurological injury, people may adopt individual- and leg-specific strategies to accelerate the CoM.

However, characterizing changes in template signatures with exoskeletons or following neurological injury requires addressing a major methodological challenge: Manually identifying optimal template signature structure is a slow, *ad hoc* process that relies on first-principles knowledge of the system^19^. Using this manual approach, comprehensively comparing candidate template signatures for even a moderate number of template mechanisms is challenging: the number of comparisons increases combinatorially with the number of candidate mechanisms. New approaches are needed to select mechanisms rapidly and systematically from a literature-based library of candidate mechanisms.

Recent advances in data-driven modeling and machine learning provide powerful tools to identify template signatures from walking data^29-32^. One such algorithm, **Hybrid-SINDy** (*SINDy: Sparse identification of nonlinear dynamics*^29^), identifies sparse nonlinear dynamics in hybrid systems from time-series data, making it particularly appropriate for identifying template models of walking, which have distinct dynamics based on foot contact configuration^28,31^. Hybrid-SINDy automatically identifies and compares a large number of candidate dynamical models (*e.g*., template signatures) from an arbitrary library of possible functional forms (*i.e*., mechanisms). The algorithm uses information criteria to determine the relative plausibility of each candidate model and selects only those that are highly plausible (i.e., that are both parsimonious and highly representative of the system)^33^. When applied to human walking data this approach will, therefore, enable rapid, systematic identification of individual-specific template signatures.

The purpose of this study was to identify changes in template-signature-based representations of CoM dynamics in response to ankle exoskeletons. We used the Hybrid-SINDy algorithm to identify physically interpretable, low-dimensional template signatures describing CoM dynamics during walking in unimpaired adults and evaluated how template signature coefficients changed with hinged and stiff ankle exoskeletons. We hypothesized that the addition of ankle exoskeleton frame and stiffness would alter template signature coefficients. Additionally, to examine the potential of Hybrid-SINDy-based template signatures to reveal changes in CoM dynamics with ankle exoskeletons for individuals with neurological injuries, we present a case study evaluating altered template signatures in one individual with post-stroke hemiparesis.

## Methods

### Data collection

We collected 3D marker trajectories using a ten-camera motion capture system (Qualisys AB, Gothenburg, SE) and GRFs using an instrumented treadmill (Bertec Inc., Columbus, USA) in twelve unimpaired adults (6M/6F; age = 23.9 ± 1.8 years; height = 1.69 ± 0.10 m; mass = 66.5 ± 11.7 kg) and one stroke survivor with lower-limb hemiparesis (*sex not disclosed*; age = 24 years; height = 1.70 m; mass = 68.0 kg). Participants walked at their self-selected steady-state speed on a fixed-speed treadmill in shoes-only and with bilateral passive ankle exoskeletons under two conditions: zero resistance to ankle flexion (*i.e*., zero stiffness; K_0_) and high dorsiflexion resistance (*i.e*., high stiffness; K_H_ = 5.1 Nm/deg; Figure 1). The order of walking conditions was randomized. A detailed description of the acclimatization protocol and data preprocessing can be found in^34^. Briefly, in a second session following a practice session, data were collected while participants walked at their self-selected speed—determined in the practice session—for six minutes per condition, including two minutes to acclimate to the treadmill before data were recorded. Only the third and fourth minutes of data were used in this study. To mitigate fatigue, the post-stroke participant walked under the same protocol, but for only three minutes per condition^35^. This study was approved by the University of Washington Institutional Review Board (#47744). The study was performed in accordance with the approved protocol and University of Washington Institutional Review Board guidelines and regulations. All participants provided written informed consent prior to participating in the study.

**Figure 1:**
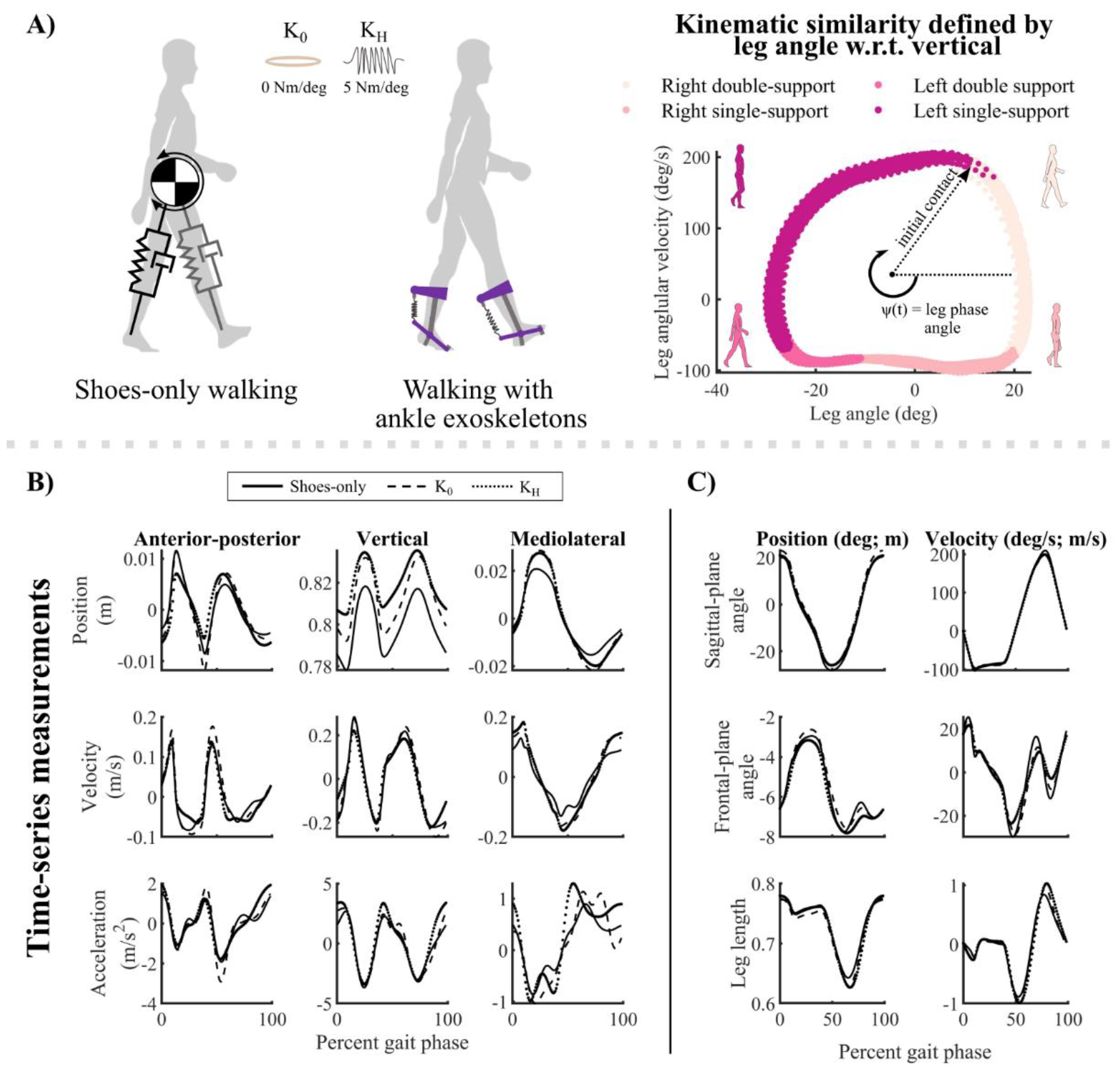
Depictions of walking conditions, phase variables, and example template state variables. A) Two-dimensional depictions of template model applied to human walking without and with ankle exoskeletons (left). The phase portrait (right) defined a phase variable, *ψ*, used to cluster kinematically similar measurements for model fitting. Colors denote gait phases corresponding to first and second double-limb support, single-limb support, and swing of the right leg. B) Stride-averaged global CoM position, velocity, and acceleration for an exemplary unimpaired adult in the anterior-posterior, vertical, and mediolateral directions. The three exoskeleton conditions are shown in panels B and C: shoes-only (solid lines), zero-stiffness exoskeletons (K_0_; dashed lines), and stiff exoskeletons (K_H_; dotted lines). C) Template position and velocity states used to fit the template signatures were defined by sagittal- and frontal-plane leg angles, and leg length.

### Estimating template signatures with Hybrid-SINDy

We used the Hybrid-SINDy algorithm to identify template signatures during walking with and without ankle exoskeletons, separately for each walking condition.

#### Kinematic variable extraction

For each exoskeleton condition, we used OpenSim’s *Body Kinematics* algorithm to estimate the CoM and foot positions^36,37^. In this section, we describe the SINDy and Hybrid-SINDy algorithms in the context of identifying template signatures, while more detailed explanations can be found in^29^ and^31^. The 3D CoM accelerations, 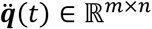 (Figure 1B), were described by continuous-time nonlinear dynamics, 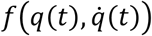, where *m* denotes the number of samples and *n* = 3 denotes the output variables:

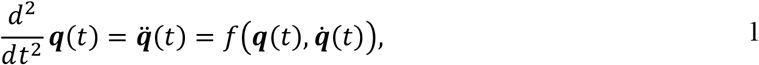

where time is denoted by *t* ∈ ℝ^*m*×1^, and *q*(*t*) and 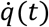 represent CoM positions and velocities relative to the feet, respectively, in ℝ^*m*×*n*^, in the anterior-posterior, vertical, and mediolateral directions. We assume that only a small number of functional forms (*i.e*., mechanisms) in 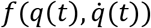 describe most of the system’s behavior. We omit the time notation in the remaining sections.

#### Sparse Identification of Nonlinear Dynamics (SINDy)

The SINDy algorithm^29^ recovers sparse nonlinear dynamics from a library of candidate functional forms, which may consist of arbitrary nonlinear functions of system measurements. Adopting the notation from^31^, we can rewrite the dynamics in equation (1) as:

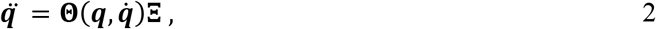

where **Ξ** ∈ ℝ^*p*×*n*^, is a linear map from nonlinear function library encoding mechanisms that may be useful in describing CoM dynamics, 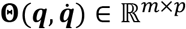, to CoM accelerations, 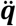. The coefficients in **Ξ**, therefore, describe how each template signature mechanism accelerated the CoM, with non-zero coefficients denoting the active terms that define the template signature structure. In the context of the template model investigated here, non-zero coefficients denote the mechanisms included in the model. We included *p* = 14 functional forms (mechanisms) in the function library (7 per leg), described below. The SINDy algorithm promotes sparsity in the model using sequential least-squares regression with hard thresholding, with the threshold defined by the sparsity parameter, *λ* (equation 3)^31^. This thresholding approach penalizes the zero-norm of **Ξ** and solves:

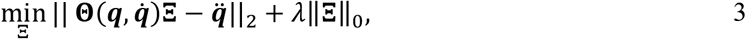

#### Hybrid-SINDy

Hybrid-SINDy extends SINDy in two important ways. First, Hybrid-SINDy uses clustering to generalize SINDy to hybrid systems. For human walking, clustering enables unique dynamics to be identified in each *gait phase*, defined by foot contact configuration (*i.e*., single- and double-limb support)^9,14,18^. We replace the term *hybrid regime*, used in the original Hybrid-SINDy manuscript, with *gait phase* for clarity in the context of human walking^31^. Second, Hybrid-SINDy uses information criteria to automatically select the system dynamics that best describe the data. This approach enables competing hypotheses about the mechanisms describing CoM acceleration to be rapidly and systematically compared, thereby highlighting mechanisms that are critical to describing CoM dynamics across individuals and mechanisms unique to a subset of individuals. Note that Hybrid-SINDy does not compare entirely distinct sets of governing equations (e.g., SLIP-like template vs. a passive dynamic walker with knees^12^). Rather, the algorithm selects which mechanisms should be included in a SLIP-like template model to best describe CoM dynamics.

### Applying Hybrid-SINDy to walking

We applied the Hybrid-SINDy algorithm to human gait using the following steps for each participant and walking condition (outlined in Figure 2; example results shown in Figure 3). Note that within each gait phase, we expanded upon the original Hybrid-SINDy algorithm by using multi-model inference to define a single template signature when multiple signatures were plausible (Step 5)^31^.

1. *Clustering* (Figure 2; Figure 3B): We used a clustering approach to increase robustness to measurement noise and identify frequently occurring template model structures^31,38^. For the first 3600 samples (30s; ∼25-30 strides) in the training set (10800 samples were available for clustering in each trial), we generated clusters of each sample’s 800 nearest neighbors and identified the centroid of each cluster (3600 clusters, total). These clusters were used to estimate template model coefficients (Step 2). During clustering, nearest neighbors were selected based on their continuous kinematic phase: the phase angle of the right/non-paretic leg angle and angular velocity relative to vertical (Figure 1A; right)^34,39^. We also used this phase variable to normalize stride progression. The last 3600 samples of each dataset were withheld from training and used during model evaluation and selection (Step 3). Some clusters contained data in both single- and double-limb support phases. However, these clusters tended to have relatively large error during model evaluation, such that they would not be selected as *plausible* in Step 3, below^31^. Further, we selected our cluster size to ensure that clusters were small enough to contain data from only one gait phase: the average cluster width (800 samples) spanned only 7.4% of the training data, smaller than the duration of double-limb support (10-12%)^40^.
2. *Model estimation* (Figure 2; Figure 3B & C): For each training cluster, we used SINDy to estimate the coefficients of multiple template signatures by sweeping 40 sparsity threshold values, ranging logarithmically from 1-100% of the largest magnitude coefficient in the full-dimensional model in each cluster. This approach typically produced 5-15 unique signatures per cluster.
3. *Model evaluation and selection*: Using the 3600 samples of held-out data, we evaluated the ability of each template signature to reconstruct CoM accelerations in the anterior-posterior, vertical, and mediolateral directions. We computed the average reconstruction error of the held-out data over the gait cycle (equation 4).

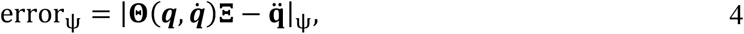

where *ψ* represents the continuous phase of the gait cycle, from 0-100% of a stride. We selected template signatures based on two criteria: First, we discarded signatures that were identified in less than 1% of training clusters. Frequently occurring template signatures are more likely to be robust to measurement noise or stride-to-stride variability, making them better representations of an individual’s gait dynamics^31^. Second, for each gait phase—single- and double-limb support—we selected the frequently occurring template signatures that had the highest likelihood according to the Akaike Information Criterion (AIC)^33,41^. The AIC is widely used to compare candidate representations of a system (*e.g*., template signatures) according to their number of free parameters and log-likelihood^33^. According to the AIC, a candidate representation that has a lower AIC score than competing representations is considered the most plausible (*i.e*., best) candidate representation of the system. Adopting the formulation in^31^, assuming that model errors are independently, identically, and normally distributed, the AIC can be written in terms of the number of free parameters, *k*, number of samples, *ρ*, and the sum of squared residuals:

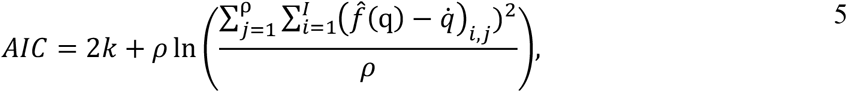

where in the outer summation is over *ρ* = 3600 samples and the inner summation, is over the *I* = 3 output states. The AIC favors parsimonious, highly representative models, which is ideal for identifying minimalist representations of gait dynamics. Like Mangan and colleagues^31^, we used the AIC corrected for finite sample sizes (AICc):

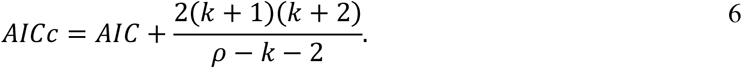

The correction term approaches zero as the number of samples, *ρ*, increases. We then determined the relative plausibility of competing template signatures using their relative AICc score, *ΔAICc*_*j*_^31,33^:

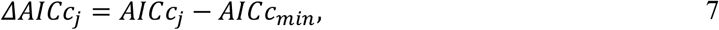

where ΔAICc_j_ represents the relative AICc score for the *j*^*th*^ model. *AICc*_*min*_ represents the AICc of the model with the lowest AICc among the models compared within a gait phase. The best model according to the relative AICc has a score of *ΔAICc*_*j*_ = 0 and all other models had higher scores. Burnham and Anderson noted that models with *ΔAICc*_*j*_ ≤ 2 have substantial support, while *ΔAICc*_*j*_ > 7 have low support^33^. We adopted the threshold of^31^, deeming template signatures with *ΔAICc*_*j*_ ≤ 3 to be *plausible*.
4. *Multi-model inference* (Figure 2): Since human gait dynamics are not strictly hybrid and template models are approximations of CoM dynamics, multiple template signature structures may be plausible in each gait phase. To construct a single template signature for each gait phase, we computed a weighted-average signature using Akaike weights, *ω*_*j*_, where *j* is the *j*^*th*^ plausible model in the gait phase^33^. Note that we performed multi-model inference separately for each gait phase. Akaike weights are defined as:

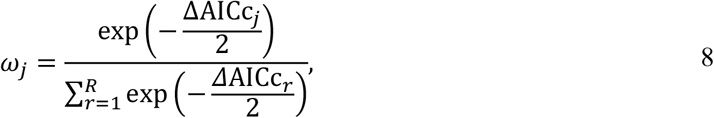

where 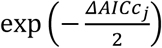 defines the likelihood of the *j*^*th*^ template signature given the observations^33^. The denominator denotes the summation of exponentially scaled relative AIC scores over all *R* candidate models. This approach weighs each signature based on its likelihood relative to the other plausible signatures.
5. *Uncertainty estimation and model accuracy* (Figure 2; Figure 3D): To evaluate the robustness of the template signatures to noise and stride-to-stride variations in the data, we performed 200 bootstrapped estimates of each template signature coefficient in each gait phase separately for each trial. Each bootstrapping iteration randomly selected 3600 samples to estimate template signature coefficients, with replacement. We quantified the robustness of each template signature coefficient to variability in the data using the coefficient of variation (CV) of each participant and condition^27^. Template signature coefficients for each participant, condition, and gait phase, were defined by the mean of the bootstrapped estimates. Figure 3D shows the estimated coefficients of the bootstrapping procedure applied to a synthetic SLIP model (*see Supplemental – S2*). Synthetic SLIP model parameters and simulation results (Figure 3D; gray bars and trajectories, respectively) were used as ground truth to validate the algorithm’s ability to identify template models of walking (*Supplemental – S2*).

**Figure 2:**
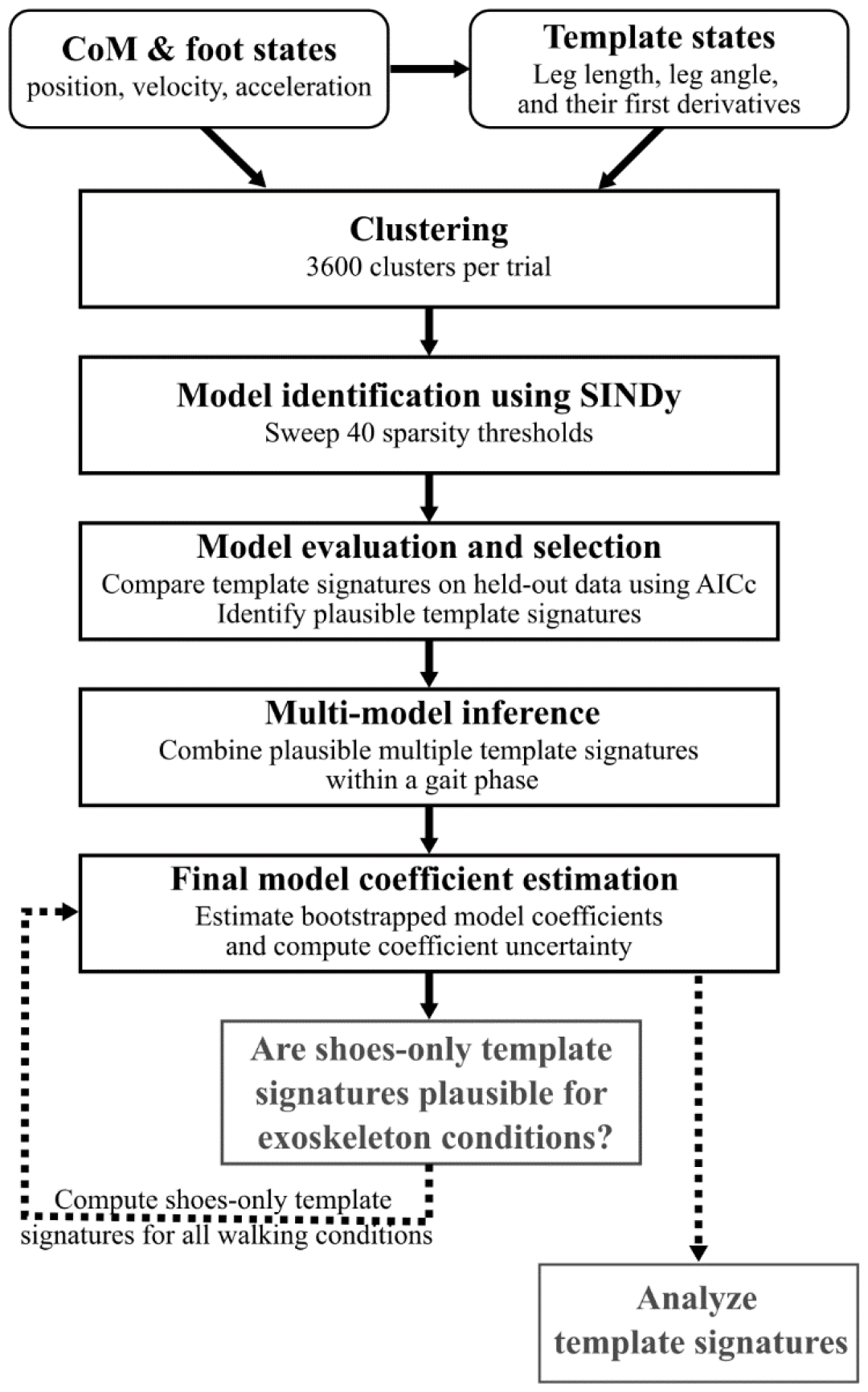
Block diagram depicting data processing pipeline to calculate template signatures for each trial. Boxes with rounded corners denote data pre-processing steps. Black boxes with square corners denote steps using the Hybrid-SINDy algorithm. Gray boxes denote analysis steps. The process starts from template state variable calculation, followed by clustering of kinematically similar samples, model estimation, and model selection. Multi-model inference defines a single template signature in each gait phase if multiple signatures are plausible. Bootstrapping is then used to estimate final model coefficients and quantify uncertainty in each coefficient. Because shoes-only template signatures were plausible for exoskeleton conditions, we repeated the model coefficient estimation step for the exoskeleton conditions using the shoes-only template signature structure (dashed arrow). The resulting template signatures were used in analysis. Full details of each step are described in the section: *Applying Hybrid-SINDy to walking*.

**Figure 3:**
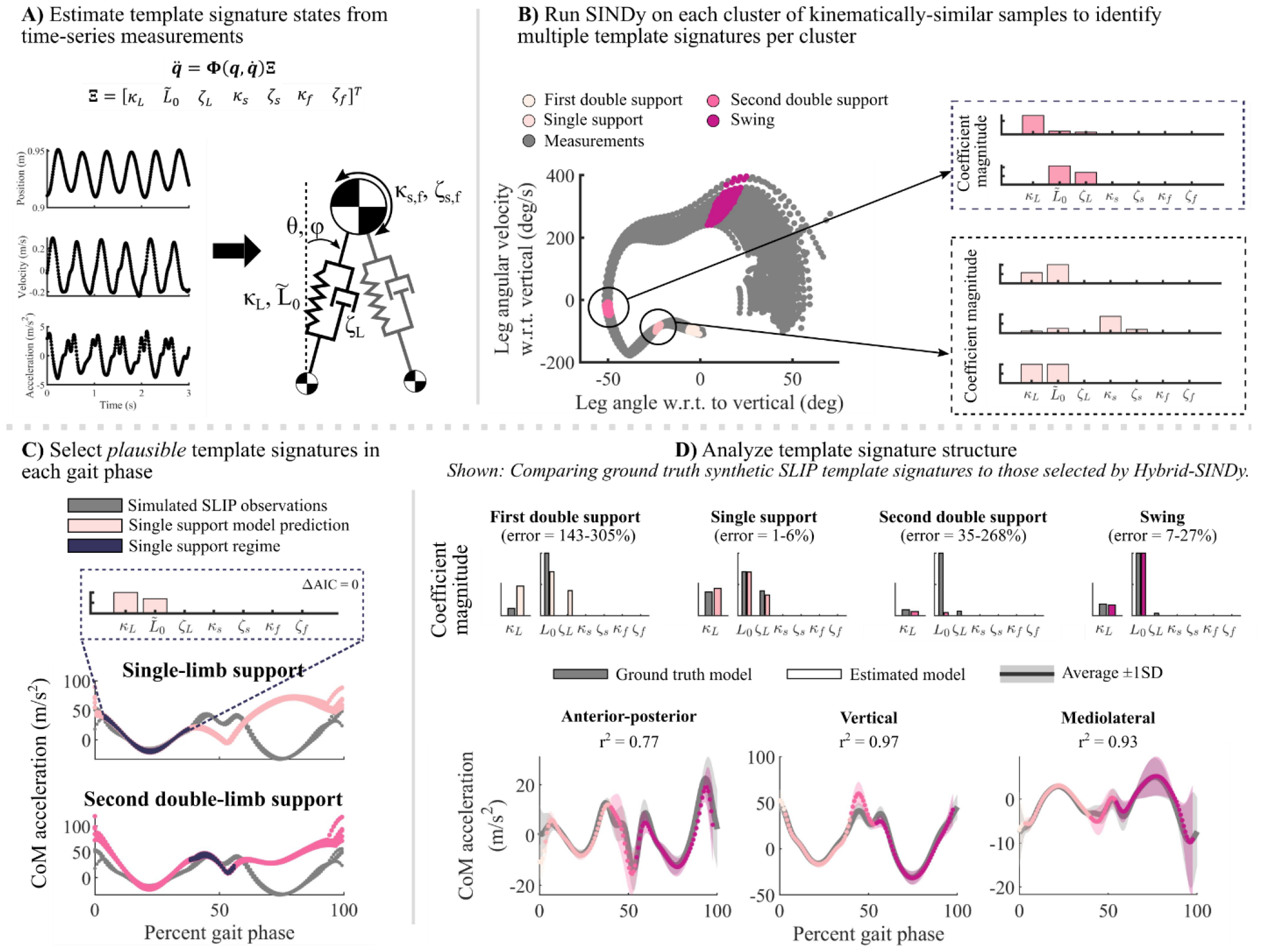
Example of the Hybrid-SINDy algorithm, as applied to a ground-truth synthetic SLIP walking model. Details of model validation using a synthetic SLIP can be found in Supplemental – S2. A) Example of simulated CoM kinematics from the synthetic SLIP and a diagram of the full-dimensional template model used to estimate template signatures. Each coefficient in the diagram corresponds to one element of the function library, **Ξ**, which maps between nonlinear Template states, 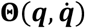 to reconstruct CoM accelerations, 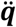. Template state variables are described in Table 1. B) Clustering of kinematically similar samples. The phase portrait (left) of simulated data from the synthetic SLIP (gray) is analogous to that shown in Figure 1A for human walking data. Colors denote clusters in different gait phases. Example clusters (colors) are shown for each gait phase. Multiple candidate template signatures are estimated for each cluster (right; examples of signatures in two clusters). C) In each gait phase, template signatures were compared using the Akaike Information Criterion (AIC). The plots show synthetic SLIP CoM accelerations (gray) and template signature reconstructions (colors) in single-limb (top) and second double-limb (bottom) support. Model errors were low only within the gait phase containing the model’s cluster centroid (purple). D) Model coefficients (top) can be compared between walking conditions. Bar plots show ground truth synthetic SLIP parameters (gray) and the corresponding estimated template signature coefficients (colors). The range of percent errors in coefficients are shown above each plot; analogously, differences in human template signatures can be compared between walking conditions. The average (±1SD) ground truth CoM accelerations from the synthetic SLIP (bottom; gray) and model predictions in each gait phase (colors). Model accuracies are quantified using the coefficient of determination (r^2^).

### Template signatures mechanisms and dynamics

To model three-dimensional CoM dynamics during walking, we created a function library of candidate mechanisms based on prior literature (Figure 3A; Table 1). We included *leg springs*^13,19-21,25^ and *dampers*^26^, which produce force along the leg. Leg springs are common energetically conservative mechanisms used to describe walking and running dynamics and enable a double-limb support phase. Leg dampers are less common, but have been used to capture non-conservative gait dynamics^26^. We also included *rotary springs*^13,19^ and *dampers* in the sagittal and frontal planes, which enable forcing transverse to the leg axis. The addition of rotary springs has been shown to improve reconstructions of anterior-posterior GRFs in a bipedal SLIP^19^. We did not identify rotary damping elements in prior literature but included them as candidate mechanisms describing non-conservative forcing transverse to the leg.

**Table 1.**
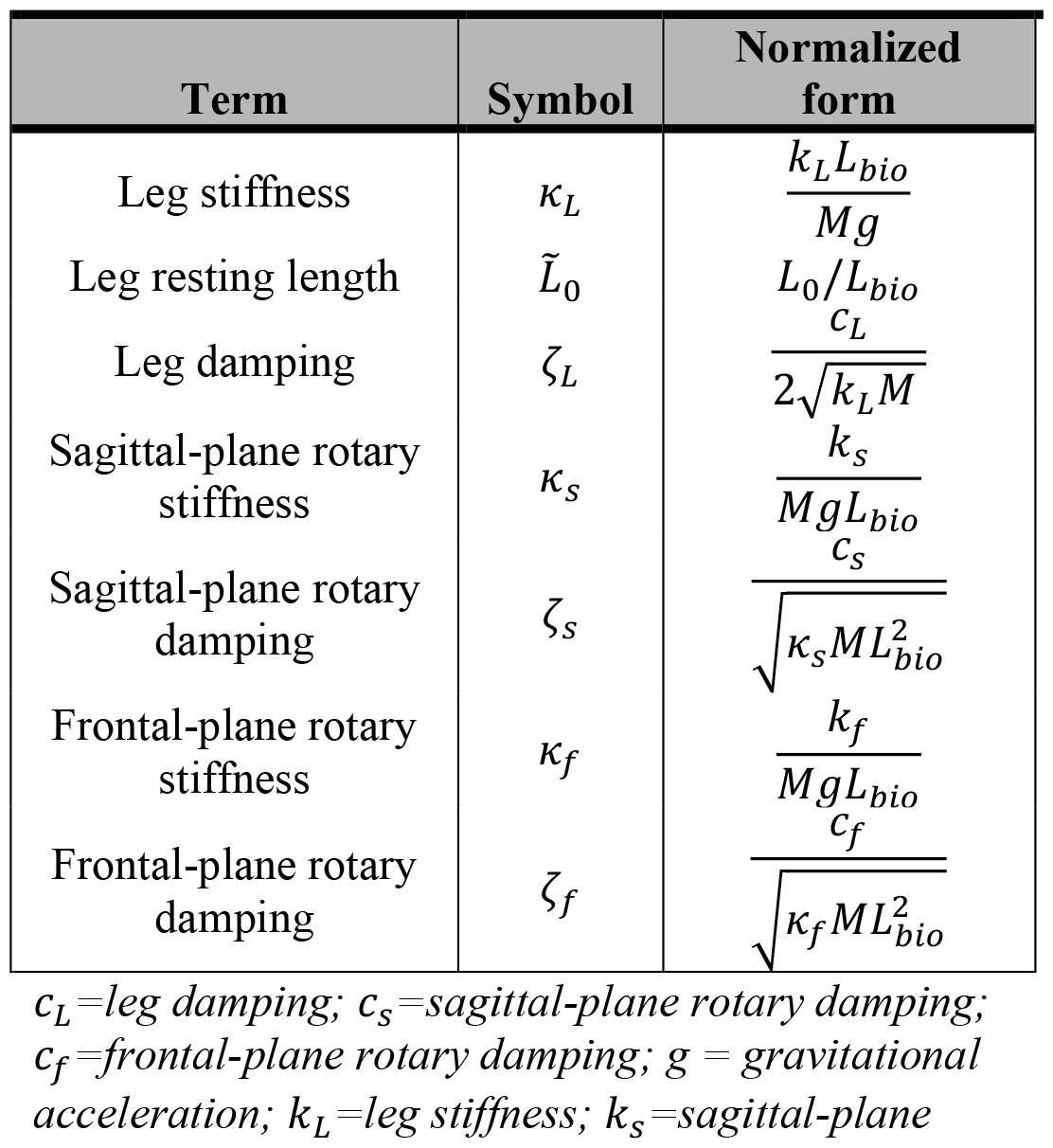
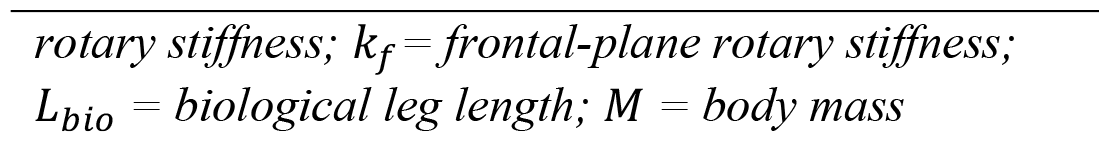
Template signature functional forms.

We included only passive mechanical elements in the mechanism library because these elements can be used in a hybrid modeling framework to approximate active control of walking, such as for lateral stabilization or to inject and dissipate energy^19,42-44^. Active mechanisms or state-based controllers could produce more parsimonious models but would be more challenging to interpret.

The mechanisms selected by the Hybrid-SINDy algorithm define the template signature *structure*, which describes characteristic strategies to accelerate the CoM. The identified template signature *coefficients* describe each mechanism’s contribution to CoM accelerations.

The dynamics of a three-dimensional bipedal SLIP augmented with damping and rotary mechanisms may be written as

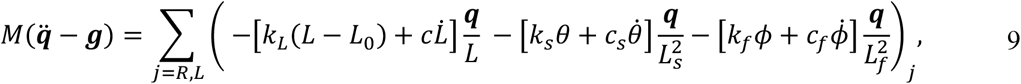

where *M* is body mass, ***g*** is the gravity vector, *ϕ* describes the traverse-plane leg angle, and *θ* describes the leg angle from vertical in the direction defined by *ϕ* (Figure 1C)^19,25^. The summation represents the total force generated by the legs on the CoM. The left-most brackets contain mechanisms that impart forces radially along the leg: *k*_*L*_ is the leg stiffness, *L* is the instantaneous leg length (Figure 1C), *L*_0_ is the leg resting length, *c*_*L*_ is the leg damping, 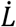 is the instantaneous leg velocity. We henceforth denote *L*_0_ as *leg length* for clarity. The middle bracket contains mechanisms that impart forces transverse to the leg axis in the sagittal plane: *k*_*s*_ and *c*_*s*_ are the sagittal-plane rotary stiffness and damping, respectively. *L*_*s*_ denotes the sagittal-plane leg projection. Analogously in the right-most brackets, *k*_*f*_ and *c*_*f*_ represent the frontal-plane rotary stiffness and damping, respectively, and *L*_*f*_ denotes the frontal-plane leg projection. The derivation of system dynamics can be found in *Supplemental – S1*.

### Normalized template mechanisms

To account for inter-individual differences in walking speed and body size during analysis, we normalized the template signatures (Table 1). Leg stiffness was normalized as in^17,19,31^. Leg resting length was normalized to the measured leg length^19,26^. Rotary stiffness was normalized according to^19^. All damping terms were converted to damping ratios^26^. The normalized leg, sagittal-plane, and frontal-plane stiffness mechanisms are denoted by *k*_*L*_, *k*_*s*_, and *k*_*f*_, respectively. The normalized leg, sagittal-plane, and frontal-plane damping mechanisms are denoted by *ζ*_*L*_, *ζ*_*s*_, and *ζ*_*f*_, respectively. Normalized leg length is denoted 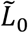. We can rewrite equation (9) as a linear combination of our normalized coefficients and nonlinear transformations of our states:

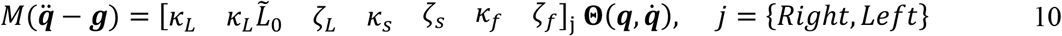

The CoM position and velocity relative to the feet were used to compute candidate template signature states: leg lengths and lengthening velocities, sagittal-plane leg angles and angular velocities relative to vertical, and frontal-plane leg angles and angular velocities relative to vertical. The complete function library can be found in *Supplemental – S1*.

### Evaluating Hybrid-SINDy’s ability to select template signatures

We evaluated the Hybrid-SINDy algorithm’s ability to accurately identify walking dynamics in the presence of noise and an incomplete mechanism library (*i.e*., missing functional forms relative to the true system dynamics) using forward simulations of a bipedal SLIP walking model (example shown in Figure 3)^25^. The synthetic SLIP model analysis and results are described in *Supplemental-S2*.

### Identifying template signatures in human gait

To quantify how well template signatures captured COM dynamics, we computed coefficients of determination (*r*^*2*^) between the measured CoM accelerations and those predicted by each participant’s template signatures, averaged over the anterior-posterior, vertical, and mediolateral directions.

To evaluate the extent to which each mechanism described CoM accelerations in unimpaired adults, we determined the proportion of participants for whom each template signature coefficient was selected and the average number of non-zero mechanisms in each gait phase. Template signature terms that are identified across individuals may represent mechanisms fundamental to CoM dynamics, while infrequently identified mechanisms may describe individual-specific features of CoM dynamics.

To determine if unimpaired CoM dynamics during shoes-only walking generalized to walking with ankle exoskeletons, we evaluated the ability of shoe-walking template signature structures to reconstruct CoM accelerations in the K_0_ and K_H_ conditions. We used least-squares regression to estimate template signature coefficients for the K_0_ and K_H_ conditions using the shoes-only template signature structure. We compared the AICc scores between these signature structures and those of the signature structures specific to the K_0_ and K_H_ trials. To determine if shoes-only template signatures were less plausible than signature structures selected for the K_0_ and K_H_ conditions, we used one-sample right-tailed t-tests (*α* = 0.05) to test if differences in the average relative AICc scores were greater than three (e.g., *AICc*_*Shoe*_ − *AICc*_*K*0_ > 3 for the K_0_ condition)^33^.

To determine if the ankle exoskeleton frame or mass impacted CoM dynamics, we compared template signature coefficients between the K_0_ and Shoe conditions. Similarly, to evaluate the impacts of exoskeleton stiffness on CoM dynamics, we compared template signature coefficients in the K_H_ and K_0_ conditions. For both comparisons, we used paired independent-samples t-tests with Holm-Sidak step-down corrections for multiple comparisons (*α* = 0.05)^45^. Because shoes-only template signature structures were plausible for most unimpaired participants, we re-estimated template signatures in the exoskeleton conditions using the shoes-only template signature structure before comparing coefficients across walking conditions.

To determine if CoM dynamics may be altered post-stroke, we computed the percent difference in the non-paretic and paretic leg template signature coefficients during shoes-only walking in one individual with post-stroke hemiparesis. We also evaluated changes in post-stroke CoM dynamics with ankle exoskeletons by computing percent changes in template signature coefficients for the K_0_ condition compared to the shoes-only and K_H_ conditions.

## Results

When walking in shoes-only, template signatures reveal common and more individual-specific representations of CoM dynamics across unimpaired participants. Unimpaired template signatures were not significantly different between legs (paired 2-sample t-test; p > 0.080). In all gait phases, SLIP mechanisms—leg stiffness and leg length—were selected (*i.e*., had non-zero coefficients) in 100% of legs (*k*_*L*_ and *L*_0_ in Figure 4). Rotary stiffness and damping mechanisms were selected in less than 30% of legs in single-limb support and swing. On average across participants and legs, 2.3 ± 0.8 and 2.9 ± 1.3 terms had non-zero coefficients in the stance and swing legs, respectively. More mechanisms were selected during double-limb support phases: Rotary stiffness terms were selected in 79-83% of legs in the leading leg and 67-79% of legs in the trailing leg. Damping mechanisms were selected most frequently (33-67%) in the double-limb support phases (*ζ* terms in Figure 4). On average, 5.0 ± 1.8 and 4.6 ± 1.6 terms had non-zero coefficients in first and second double-limb support, respectively.

**Figure 4:**
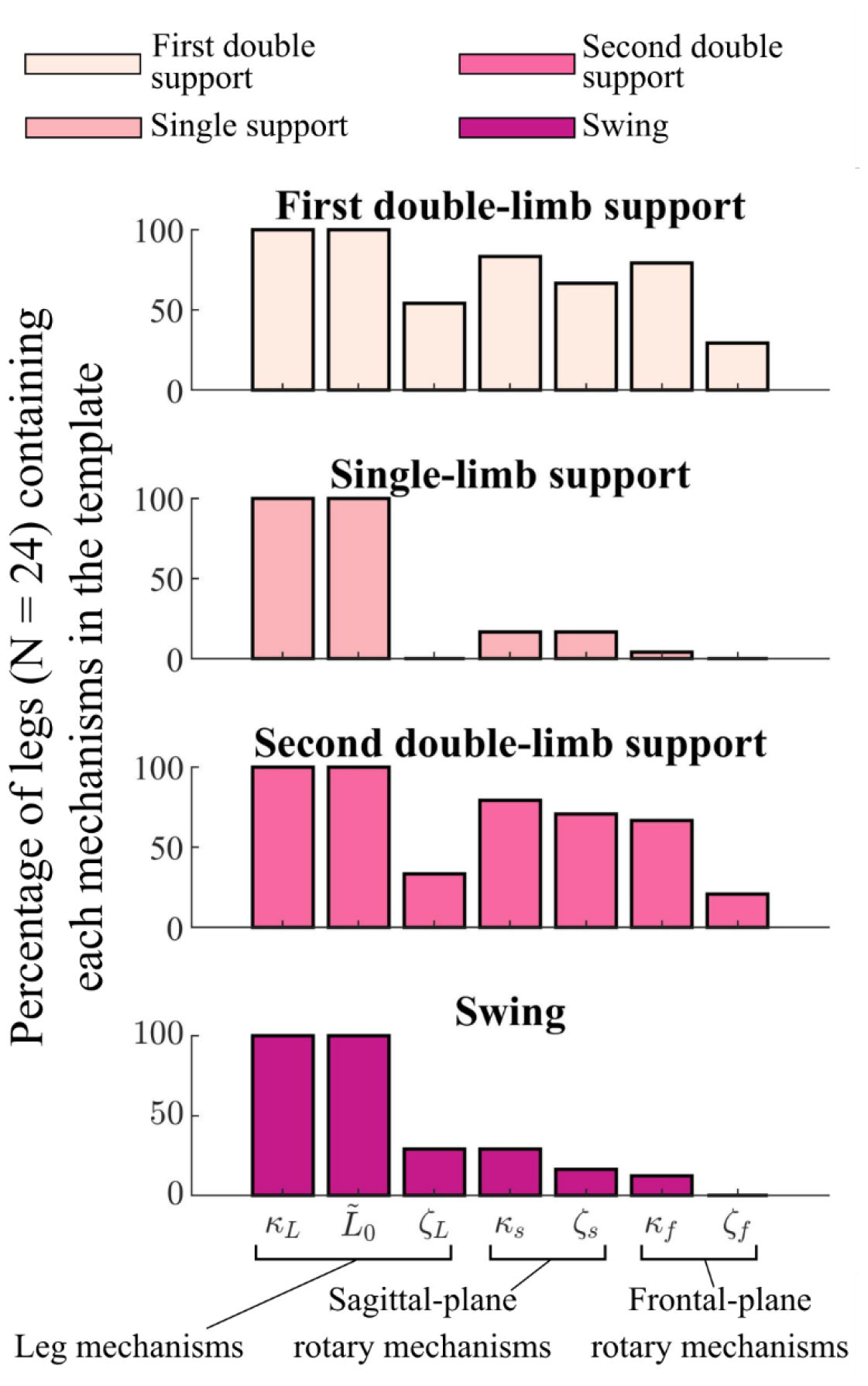
The percentage of unimpaired legs whose template signatures contained each mechanism in each gait phase. For 24 legs, the percentage of legs for which each template signature mechanism was selected by the Hybrid-SINDy algorithm. Colors denote each gait phase. Mechanisms selected in a larger percentage of legs suggest common representations of CoM dynamics, while less frequently selected mechanisms reflect more individual-specific template features describing CoM dynamics.

For each unimpaired participant, shoes-only template signature coefficients were reliable, having low bootstrapped coefficients of variation (CVs), during single-limb support and swing: stiffness and leg length CVs were less than 0.02 and 0.06, respectively (Figure 5A). In both double-limb support phases, coefficient estimates were less reliable: CVs ranged from 0.06-4.34 across coefficients. During swing, coefficient estimates were generally reliable (CV = 0.03-0.10).

**Figure 5:**
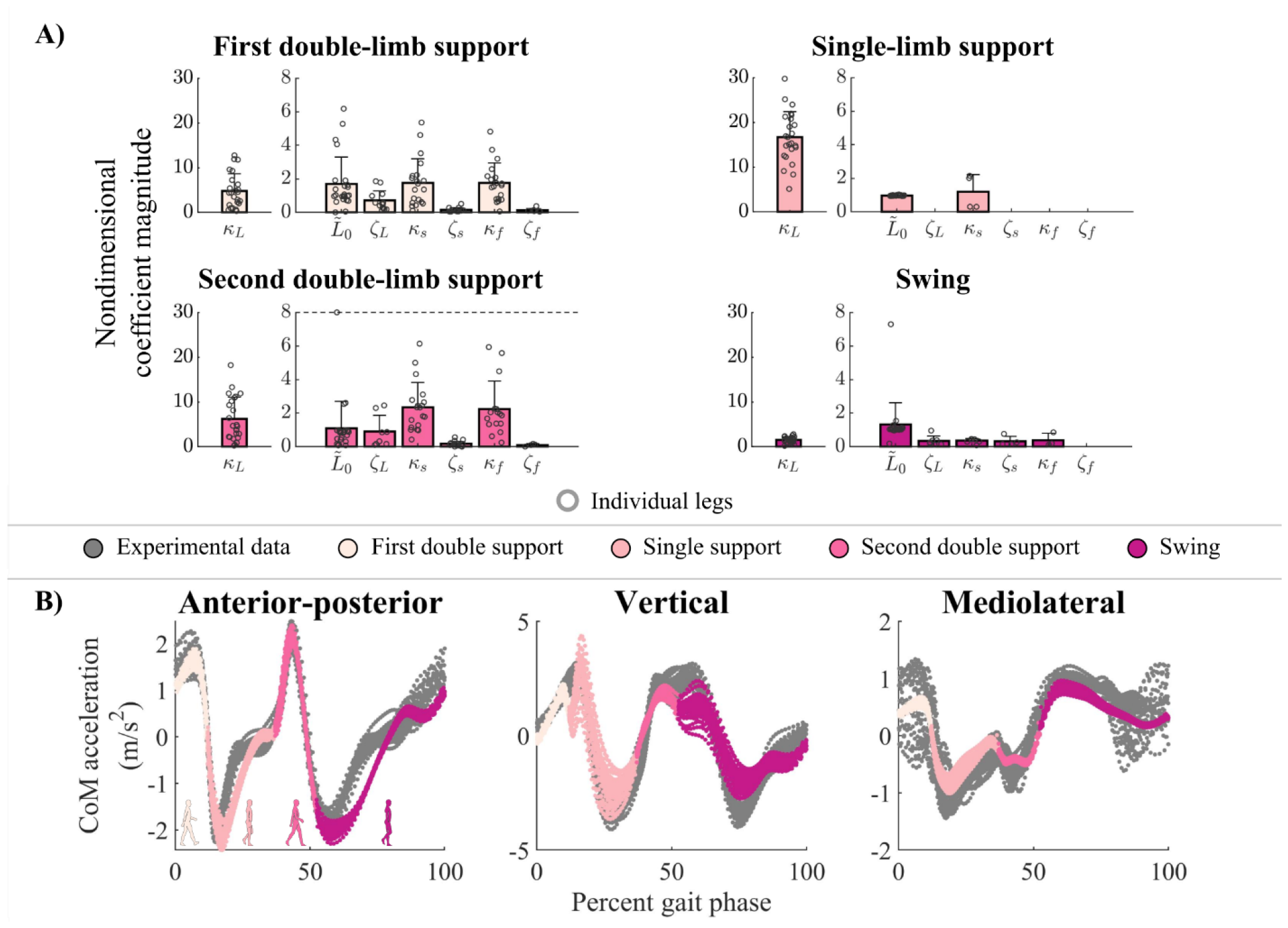
Unimpaired template signatures of walking in shoes-only. Normalized template signatures (top) and reconstructed CoM accelerations (bottom) for shoes-only walking in each gait phase. A) Bars denote the average template signature (+1SD) in single- and double-limb support. For unimpaired adults, the left and right leg coefficient estimates were grouped in each gait phase (small circles; up to 24 samples per bar). The single-limb support and swing plots contain coefficient estimates from single-limb support or swing of each leg, respectively. The first and second double-limb support phases show the coefficients of the leading/trailing leg, respectively. Note that we omitted mechanisms that were selected in less than 25% of participants for clarity. The dashed lines truncate large terms for clarity. B): Experimental (gray) and reconstructed (colors) CoM accelerations from the test dataset of an example unimpaired participant.

Across unimpaired participants, single-limb support leg length was the least variable coefficient (0.97 ± 0.03), while leg stiffness values were more variable between participants (16.7 ± 5.8; Figure 5A). Interindividual variability in double-limb support leg and rotary stiffness was larger than single-limb support coefficients. For example, leg stiffness (1.7 ± 6.0 in first double-limb support) was lower and more variable than in single-limb support. During swing, leg stiffness was relatively small (-1.5 ± 0.8) compared to single-limb support. Template signatures explained 83 ± 7% (range: 67-94%) of the variance in participants’ CoM (Example reconstruction of experimental data in Figure 5B).

Passive ankle exoskeletons elicited only small changes in unimpaired CoM dynamics: in single-limb support, the shoes-only template signature structures reconstructed CoM dynamics with similar accuracy to template signature structures selected specifically for the K_0_ condition (mean difference in *AICc* = −1.5 ± 8.4; p = 0.99) and the K_H_ condition (mean difference in *AICc* = 6.8 ± 14.6; p = 0.10; Figure 6A). Negative AICc scores indicate that the shoes-only template signature structure was more plausible than the exoskeleton-specific signature structure, which can occur if the shoes-only template signature structure is not identified in individual clusters for the exoskeleton conditions. In double-limb support, shoes-only signatures were not statistically less plausible than K_0_ or K_H_ CoM template signatures (p > 0.657). However, the relative AICc scores were highly variable, ranging from -258 ≤ Δ*AICc* < 282, such that double-limb support did not reliably indicate the plausibility of shoes-only template signature structures for the exoskeleton conditions. Because shoes-only template signatures were reliably plausible for gait with exoskeletons in single-limb support, we constrained each participant’s template signature structures to their shoes-only signature structure. Therefore, we compared template signature coefficients between exoskeleton conditions using coefficients fit to each participant’s shoes-only signature structures.

**Figure 6:**
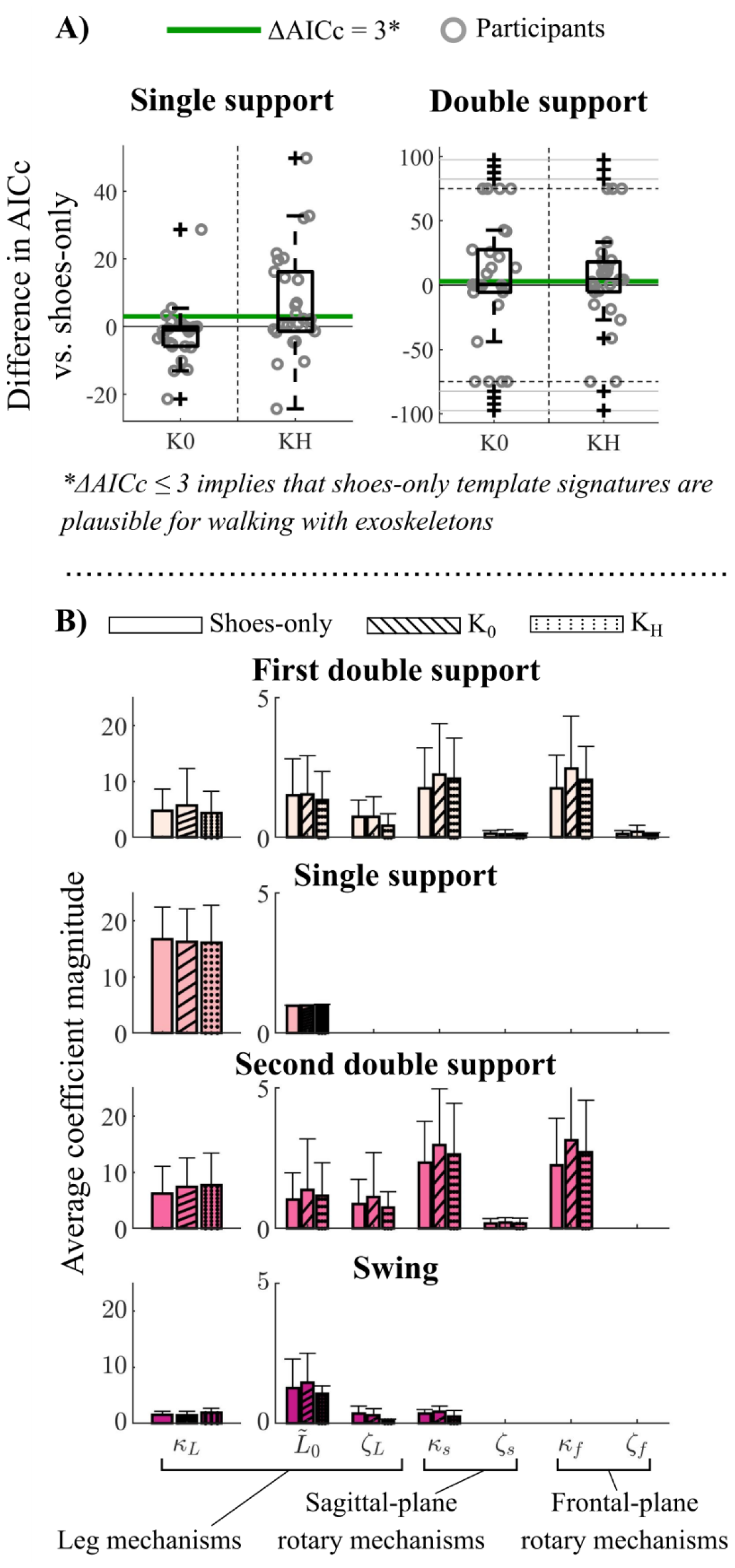
Changes in template signatures with ankle exoskeletons. A) Differences in AICc scores between template signatures identified specifically for the K_0_ and K_H_ conditions, and signatures constrained to the shoes-only signature structures. Positive Δ*AICc* values indicate that shoes-only signature structures were less plausible than those identified specifically for each condition. Δ*AICc* scores are shown for single-limb support and swing (left) and double-limb support (right). Δ*AICc* ≤ 3 (green lines) denote that shoe template signature structures were plausible for the exoskeleton conditions. For double-limb support, large Δ*AICc* scores are truncated at Δ*AICc* = ±75 for clarity. B) Template signatures of walking in shoes only (solid bars), zero-stiffness (*K*_0_; slashed bars) exoskeletons, and high-stiffness exoskeletons (*K*_*H*_; dotted bars) during single- and double-limb support. Bars represent the average (+1SD) template signature across participants. Leg stiffness (*κ*_*L*_) is on a separate subplot for clarity.

Using this approach, we found that the only significant difference in unimpaired template signature coefficients was in the leg length coefficient during single-limb support, which differed between walking in shoes-only and the zero-stiffness exoskeletons (K_0_; p = 7.9e-4; α_Sidak_ = 9.2e-4). However, this change was small 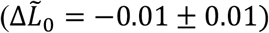, with leg length being slightly longer in the shoes-only condition. Neither ankle exoskeleton mass and frame (K_0_) nor stiffness (K_H_) altered other template signature coefficients in any gait phase (p > 0.16; α_Sidak_ = 9.3e-4) (Figure 6B).

One stroke survivor’s shoes-only template signature was symmetric in single-limb support, swing, and first double-limb support, but was asymmetric in second double-limb support (Figure 7). In single-limb support and swing, leg stiffness and leg length mechanisms were selected and reliably estimated for both legs, with sagittal-plane rotary stiffness also selected in swing (all CV < 0.02). In single-limb support the paretic leg (*κ*_*L*_ = 11.0; white bars in Figure 7) was slightly (6%) stiffer than the non-paretic leg (*k*_*L*_ = 10.4; colored bars in Figure 7). During first double-limb support, template signature coefficients differed between legs, but only the paretic leg coefficients were reliably estimated (CV = 0.00-0.06). Conversely, in second double-limb support, the sagittal- and frontal-plane rotary stiffness mechanisms were selected for the paretic, but not the non-paretic leg. Unlike first double-limb support, non-paretic leg stiffness and resting length coefficients were more reliable (CV = 0.00-0.26) than those in the paretic leg (CV = 0.34-8.01).

**Figure 7:**
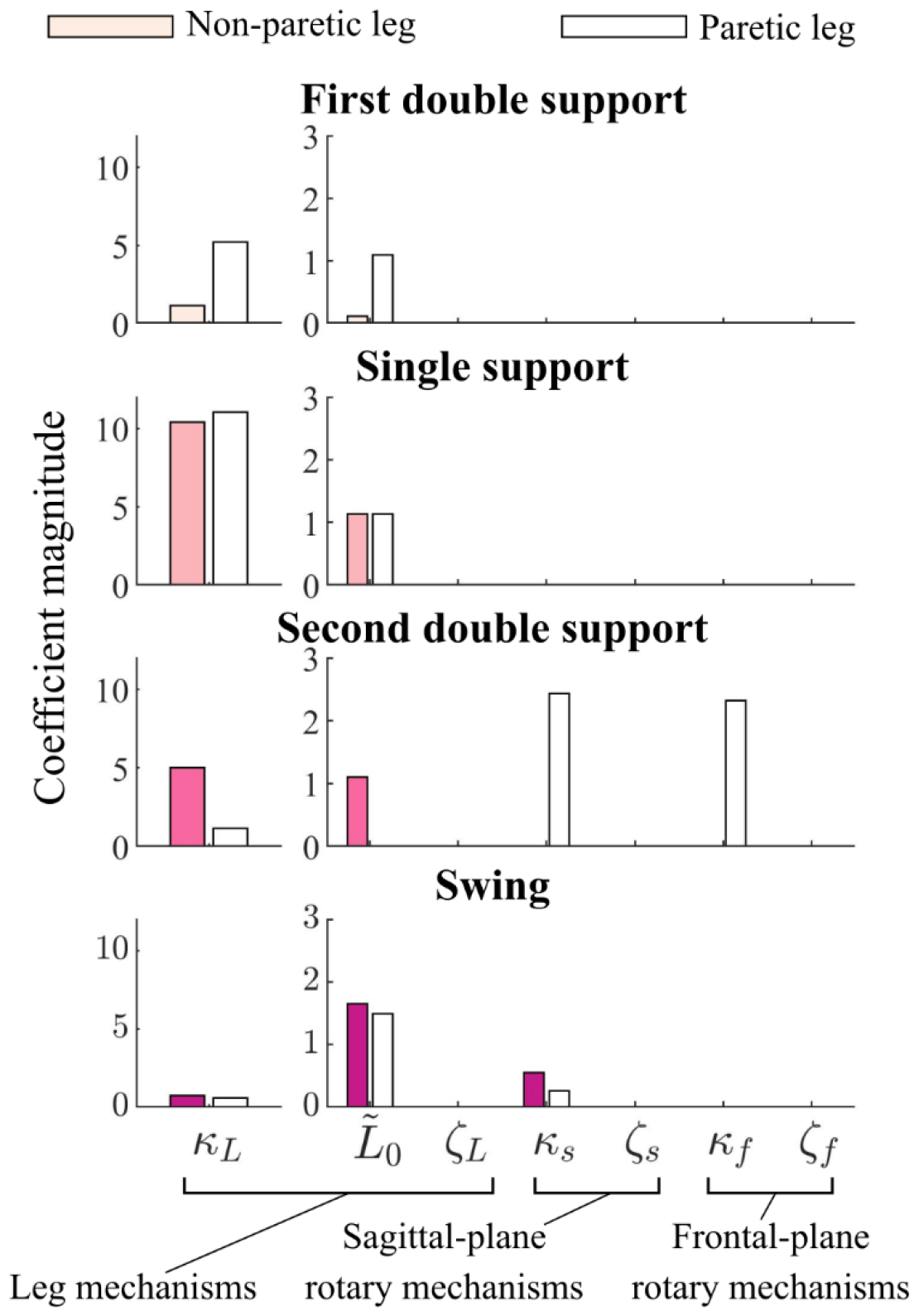
Inter-leg differences in template signatures in one stroke survivor. Non-paretic (colored bars) and paretic (white bars) template signatures for one individual with post-stroke hemiparesis. Each plot represents a different gait phase.

The exoskeleton mass and frame (Shoe vs. K_0_) primarily impacted the stroke survivor’s non-paretic and paretic leg stiffness in single-limb support, and paretic leg rotary stiffness in second double-limb support (Figure 8). Note that these coefficients were reliably estimated by Hybrid-SINDy. In single-limb support, the zero-stiffness (K_0_) exoskeleton template signatures had 33% greater leg stiffness in the paretic leg and 19% lower leg stiffness in the non-paretic leg compared to shoes-only signatures (Figure 8; slashed bars vs. no bars). In second double-limb support, sagittal and frontal plane rotary stiffness were 37 and 50% greater, respectively, in the paretic limb in the K_0_ condition compared to shoes-only. Exoskeleton stiffness (K_0_ vs. K_H_) had smaller impacts on paretic leg template signatures: in single-limb support, paretic leg stiffness was 22% less than in the K_0_ condition (Figure 8; dotted vs, slashed bars). Conversely, non-paretic leg stiffness was 85% lower in double-limb support. Paretic leg sagittal- and frontal-plane rotary stiffness were both 33% lower in second double-limb support than in the K_0_ condition.

**Figure 8:**
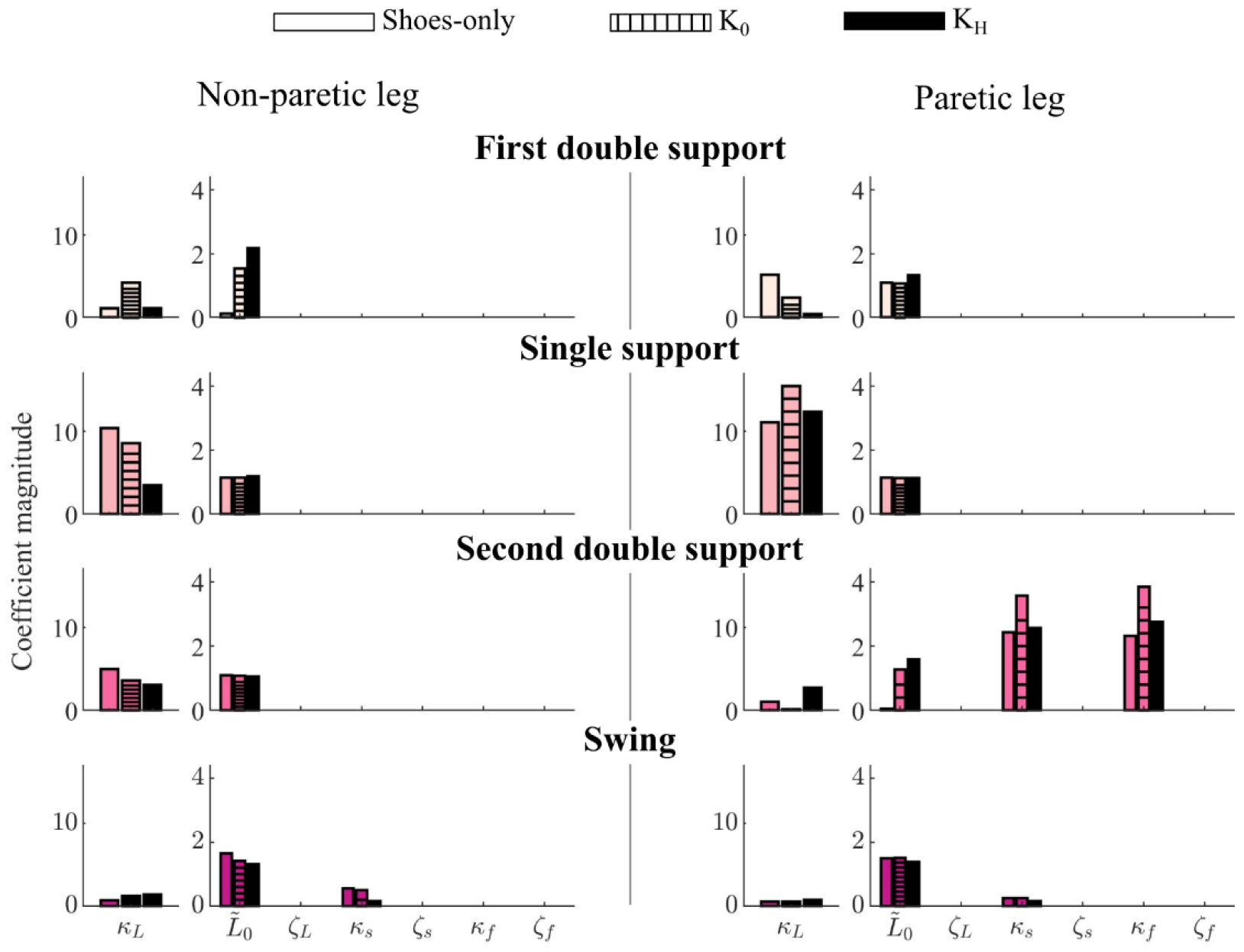
Template signatures of one stroke survivor walking with and without ankle exoskeletons. Bars denote template signatures in the shoes-only (solid color bars), zero-stiffness (*K*_0_; dashed bars), and high-stiffness (*K*_*H*_; solid black bars) ankle exoskeleton conditions for the non-paretic (left) and paretic (right) legs. Bars represent each leg’s average template signatures over 200 bootstrapped model fitting iterations. Colors correspond to each gait phase.

## Discussion

We evaluated the impacts of passive ankle exoskeletons on individual-specific template-based representations of CoM dynamics—described by template signatures—using a recently developed data-driven modeling framework, Hybrid-SINDy. Despite balancing model accuracy with parsimony, template signatures captured CoM dynamics with similar accuracy to prior work using pre-defined 2D template structures^20,26^. The symmetric and SLIP-like unimpaired template signatures automatically selected by Hybrid-SINDy during walking in shoes-only were consistent with prior template models of CoM dynamics during walking, but suggest that dynamics described by rotary mechanisms represent more individual-specific structures describing CoM accelerations^19,25^. Contrary to our hypothesis, template-based representations of unimpaired CoM dynamics were robust to the mechanical constraints of passive ankle exoskeletons: frame and mass (K_0_), and dorsiflexion stiffness (K_H_). Conversely, in our post-stroke case study, asymmetric shoes-only signatures and changes in template signatures with exoskeletons support Hybrid-SINDy’s potential to identify interpretable representations of pathological CoM dynamics, motivating future investigation into how neurological injuries impact template-based representations of CoM dynamics with ankle exoskeletons^28^.

Unimpaired and post-stroke template signatures highlight potential inter-individual differences in CoM dynamics. The selection of leg stiffness and resting length as active mechanisms (*i.e*., terms with non-zero coefficients) in 100% of legs and their selection as the only mechanisms in single-limb support for most participants is consistent with common template walking models and supports the perspective that elastic legs are foundational mechanisms for describing CoM accelerations during unimpaired walking^12,20,21,25,46^. Conversely, our finding that rotary mechanisms were selected to describe CoM dynamics in only 33-83% of legs is consistent with their less frequent application in template walking models^13,19^. One interpretation of individual differences in selected mechanisms is that leg stiffness and resting length describe coordination patterns necessary for stable or efficient walking, while rotary mechanisms describe coordination patterns that have more individual-specific impacts on gait^47^. Alternatively, differences in the selected mechanisms may be due to covariation among template signature state variables. We observed moderate covariation between some variables, particularly in double-limb support (*Supplemental – S2: Covariation of template signature state variables*). However, Hybrid-SINDy penalizes the selection of strongly covarying states, such that both variables would not likely be selected unless they independently increase model likelihood^31,41^.

Contrary to our primary hypothesis, unimpaired CoM dynamics are robust to altered ankle constraints due to passive ankle exoskeletons, despite observed changes in kinematics and muscle activity^34^. Similarly, Collins and colleagues (2015) observed small changes in total CoM power with passive ankle exoskeletons compared to walking in shoes-only^5^. Our findings suggest that, if changes in CoM dynamics in single-limb support and swing are captured by the template-based mechanisms in our function library, these small changes in CoM power are driven by changes in CoM kinematics, rather than changes in the underlying CoM dynamics. Note that these findings may not generalize to powered ankle exoskeletons, which elicit larger changes in gait kinematics and kinetics than did our passive exoskeletons and may yield larger changes in template-based representations of CoM dynamics^2,6,34^. Note that we cannot compare coefficient estimates in those gait phases: template signatures in double-limb support were not reliably estimated. More data may be needed to robustly estimate model coefficients in shorter gait phases.

However, changes—or a lack thereof—in template signature coefficients may be biased by an incomplete function library. Because template models are incomplete representations of gait, they may be sensitive to both measurement noise and unmodeled dynamics. Our analysis of a synthetic SLIP (*Supplemental – S2: Effects of measurement noise on algorithm performance*) suggests that Hybrid-SINDy can accurately identify template signature coefficients at measurement noise levels comparable to marker-based motion capture. However, changes in true CoM dynamics with exoskeletons are not completely represented by our mechanism library: template signatures accounted for less than 94% of the variance in human CoM accelerations. For example, our mechanism library did not include torso dynamics, which are known to contribute to angular momentum regulation during post-stroke gait and may be altered with exoskeletons^48,49^. Omitting functional forms from the mechanism library induces at least 5-10% differences in template signature coefficient estimates (*Supplemental – S2: Effects of missing physics on algorithm performance*). While we encoded common functional forms from literature, larger function libraries or novel functional forms may increase the robustness of inter- or intra-individual differences in template signatures to variations in kinematics between trials^13,19-21,25,26,50^. However, even if a complete set of mechanisms is included in the library, Hybrid-SINDy may not select small but important mechanisms needed to reconstruct CoM dynamics across tasks^31,41^. Future studies should consider the tradeoff between the improved interpretability of more-parsimonious models with decreased model accuracy across the tasks of interest.

In our case study of one stroke survivor, asymmetric template signatures suggest that inter-leg differences in the mechanisms describing CoM dynamics can be automatically identified from data. Consistent with template-based studies in children with cerebral palsy, the post-stroke participant’s paretic leg was slightly (6%) stiffer than the nonparetic leg in single-limb support, and rotary stiffness mechanisms were selected only in the non-paretic leg in double-limb support^27,28^. These mechanisms may reflect a more rigid paretic leg or reliance on proximal muscles for propulsion^51,52^. However, as discussed above, the small difference in stiffness may be driven by unmodeled torso dynamics^48^. While only a case study, these results highlight the potential interpretability of subject-specific template signatures to understand how the legs contribute differentially to CoM accelerations following neurological injury.

Changes in our post-stroke case study’s template signatures with exoskeletons support Hybrid-SINDy’s ability to identify interpretable impacts of ankle exoskeletons on template-based representations of CoM dynamics. For example, increases in paretic leg stiffness (K_0_ vs. shoes-only) may stem from the exoskeleton frame restricting inversion of the paretic ankle, which the participant noted during data collection. Conversely, reduced non-paretic leg stiffness and paretic leg rotary stiffness with stiff exoskeletons (K_H_ vs. K_0_) may reflect a compensatory strategy to avoid reliance on the paretic leg, if it did not adapt effectively to the stiff exoskeleton^2^. Note that these findings represent a proof-of-concept that will not generalize across stroke survivors. Larger studies are needed to understand how individual-specific neural or biomechanical constraints may alter exoskeleton impacts on CoM dynamics^2,5,7,28,34,53^.

The Hybrid-SINDy algorithm^31,32^ was essential to discovering individual-specific and leg-specific changes in CoM dynamics with ankle exoskeletons in the present study. Conversely, manually testing all possible template signatures for our fourteen-dimensional mechanism library would require a combinatorially large number of models to be fit and compared. Conversely, Hybrid-SINDy automatically selected mechanistic representations of CoM dynamics that are consistent with the literature in a fraction of the time required to manually compare each candidate model^13,19-21,25,26^. While Hybrid-SINDy has largely been applied to synthetic systems, this work supports its ability to capture dynamics from human gait data^31,32,54^.

While alternative modeling approaches, such as principle components analysis (PCA) or non-negative matrix factorization (NMF), could identify low-dimensional representations of CoM dynamics, Hybrid-SINDy provides immediately interpretable insight into the structure of these dynamics^55,56^. Hybrid-SINDy facilitates the interpretation of CoM dynamics by selecting a sparse set of readily interpretable template variables derived from expert knowledge of human gait^12,13,19-21,25,26^. PCA or NMF-based template signatures would be harder to interpret, as they would not be sparse in the space of the template variables: each mode of PCA or NMF would likely contain non-zero coefficients for all state variables^13,19-21,25,26^. More-similar approaches to ours used stepwise regression to model ankle quasi-stiffness using mechanics-based functional forms^57^ or multi-layer optimization to identify human-like template dynamics for robot controllers^58^. Hybrid-SINDy is distinct from these approaches in its use of clustering and the AIC to evaluate the relative plausibility of competing representations of CoM dynamics, a limitation of stepwise regression^41,59,60^. For all participants, Hybrid-SINDy identified a single plausible shoes-only template signature structure in each gait phase, providing confidence that the identified signatures have strong statistical support compared to alternative signatures (*see Supplemental - S1: Comparison to alternative modeling frameworks*).

Additional limitations should constrain the interpretation of template signatures. First, human gait dynamics are continuously phase-varying rather than hybrid, such that template signatures may vary continuously over the gait cycle. Our preliminary analyses found that predicted CoM dynamics became less accurate near the transitions between single and double-limb support. Further, our prior work shows that phase-varying models of gait have higher predictive accuracy, but quantifying changes in the coefficients of continuously phase-varying models is challenging^34,50^. Because our goal was to quantify changes in dynamics rather than predict CoM motion with maximal accuracy, defining gait phases based on contact configuration was reasonable and consistent with existing hybrid template models of walking^19,20,25,31^. Second, we assumed that CoM dynamics were time-invariant, though they may change with adaptation to exoskeletons^61^. We included a two-minute adaptation period for each walking condition, but additional adaptation may elicit larger changes in participants’ template signatures. Identifying the plausibility of template signatures across or within trials could improve our understanding of how template-based representations of CoM dynamics change during adaptation^26^. Third, our test set— 90-120s of each trial recording—contained 25-30 strides of data per trial and was not a rigorous evaluation of model generalizability. Our approach identifies template signatures specific to a task and walking condition, and we do not expect these signatures to generalize across tasks or to overground walking. However, because template signature coefficients are known to vary with speed^26^ and kinematics (*Supplemental – S2: Effects of missing physics on algorithm performance*), testing on withheld speeds or walking conditions was not practical. Finally, our limited sample size may have masked exoskeleton impacts on unimpaired gait.

## Conclusions

We quantified changes in individual-specific template model-based representations of CoM dynamics in response to passive ankle exoskeletons using an interpretable physics-informed data-driven modeling framework, Hybrid-SINDy. The template mechanisms describing salient features of unimpaired CoM dynamics were insensitive to ankle exoskeleton frame or stiffness. Interpretable ankle exoskeleton impacts on template representations of CoM dynamics in a case study of one individual post-stroke support the utility of template signatures in quantifying CoM dynamics with exoskeletons in people with neurological injuries. These findings also support the potential of data-driven frameworks like Hybrid-SINDy to accelerate the investigation of individual-specific representations of CoM dynamics during walking.

## Data Availability

All experimental data and modeling code used in this study are freely available at https://simtk.org/projects/ankleexopred.

## Author contributions

M.C.R., K.M.S. and J.L.P. contributed equally to the conception and design of this work and to the interpretation of results. M.C.R. collected data, conducted all analyses, created figures, and wrote the initial manuscript draft. J.L.P. provided theoretical guidance on the implementation of the Hybrid-SINDy algorithm. K.M.S. provided experimental resources and guidance on the analysis of gait with exoskeletons. All authors contributed to substantive revisions of the manuscript.

