## Supplemental - S1 for "Quantifying changes in individual-specific template-based representations of center-of-mass dynamics during walking with ankle exoskeletons using Hybrid-SINDy"

#### Overview

This supplemental provides details of the spring-loaded inverted pendulum (SLIP) dynamics used in this work, the theory underlying Hybrid Sparse identification of nonlinear dynamics (Hybrid-SINDy), the Akaike Information Criterion (AIC), and an overview of the synthetic SLIP simulations.

1. Derivation of SLIP dynamics
2. The Akaike Information Criterion
3. Comparison to alternative modeling frameworks

### SLIP dynamics

SLIP parameters are described in Figure S1.

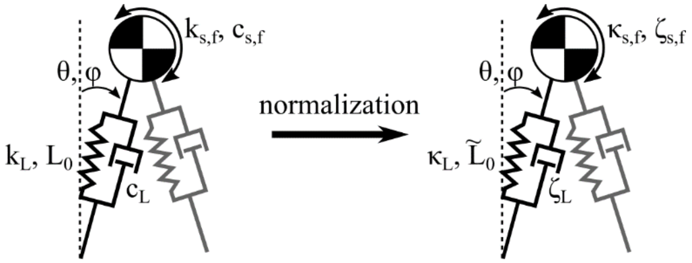


**Figure S1: A 2D schematic of the 3D SLIP with model parameters (left) and normalized parameters (right).** Variables $\theta$ and $\phi$ denote sagittal and frontal-plane leg angles with respect to vertical, respectively. The center-of-mass (COM) has mass *M*. Left: Before normalization, model parameters are denoted as: Leg radial stiffness ($k_L$) and damping ($c_L$), leg resting length ($L_0$), rotary stiffness in the sagittal ($k_s$) and frontal ($k_f$) planes, and rotary damping in the sagittal ($c_s$) and frontal ($c_f$) planes. Non-normalized parameters were estimated for each participant. Right: After normalization, parameters are denoted: Leg radial stiffness ($\kappa_L$) and damping ($\zeta_L$), leg resting length ($\tilde{L}_0$), rotary stiffness in the sagittal ($\kappa_s$) and frontal ($\kappa_f$) planes, and rotary damping in the sagittal ($\zeta_s$) and frontal ($\zeta_f$) planes[1-4]. Normalization was applied to compare participants after model fitting.

We solve the 3D SLIP dynamics with respect to linear accelerations as follows. Let SLIP model mass be denoted by *M*, and 3D positions of the center-of-mass (COM) relative to the feet be denoted *x*, *y*, & z. We solve for linear accelerations of the COM for each foot on the ground as:

$$
\begin{aligned}
\ddot{x} &= L\ddot{\theta}c\theta - L\dot{\theta}^2 s\theta + 2\dot{L}\dot{\theta}c\theta + \ddot{L}s\theta \\
\ddot{y} &= -L\ddot{\theta}s\theta - L\dot{\theta}^2 c\theta - 2\dot{L}\dot{\theta}s\theta + \ddot{L}c\theta \\
\ddot{z} &= L\ddot{\phi}c\phi - L\dot{\phi}^2 s\phi + 2\dot{L}\dot{\phi}c\phi + \ddot{L}s\phi
\end{aligned} \qquad 1
$$

where single dots denote first derivatives and double dots denote second derivatives. Other variables are defined as in Figure S1. Note that we assume that anterior-posterior and vertical dynamics are decoupled from frontal-plane dynamics. This simplification is supported by prior studies of COM dynamics during gait[5,6]. We solve radial ($\ddot{L}$) and rotary accelerations ($\ddot{\theta}, \ddot{\phi}$) as a function of spring and damper forcing terms and gravity, *g*:

$$
\begin{aligned}
\ddot{L} &= \frac{1}{M}\left[-k_L(L - L_0) - c_L\dot{L}\right] - g\cos(\theta), \\
\ddot{\theta} &= \frac{1}{ML^2}\left[-k_s\theta - c_s\dot{\theta}\right] + \frac{g}{L}\sin(\theta), \\
\ddot{\phi} &= \frac{1}{ML^2}\left[-k_f\phi - c_f\dot{\phi}\right] + \frac{g}{L}\sin(\phi),
\end{aligned} \qquad 2
$$

where variables correspond to leg radial stiffness ($k_L$) and damping ($c_L$), leg resting length ($L_0$), rotary stiffness in the sagittal ($k_s$) and frontal ($k_f$) planes, and rotary damping in the sagittal ($c_s$) and frontal ($c_f$) planes. Note that we assume that the resting angle of the rotary stiffness mechanisms is zero[2]. Substituting forcing terms into equation (1) yields:

$$\ddot{x} = \left(\frac{1}{ML^2}\left[-k_s\theta y - c_s\dot{\theta}y\right]\right) - \dot{\theta}^2 x + 2\dot{L}\dot{\theta}\frac{y}{L} + \left(\frac{1}{M}\left[-k_L(L-L_0)\frac{x}{L} - c_L\dot{L}\frac{x}{L}\right]\right)$$

$$\ddot{y} = -\left(\frac{1}{ML^2}\left[-k_s(\theta)x - c_s\dot{\theta}x\right]\right) - \dot{\theta}^2 y - 2\dot{L}\dot{\theta}\frac{x}{L} + \left(\frac{1}{M}\left[-k_L(L-L_0)\frac{y}{L} - c_L\dot{L}\frac{y}{L}\right] - g\right) \qquad 3$$

$$\ddot{z} = \left(\frac{1}{ML^2}\left[-k_f(\phi)y - c_f\dot{\phi}y\right]\right) - \dot{\phi}^2 z + 2\dot{L}\dot{\phi}\frac{y}{L} + \left(\frac{1}{M}\left[-k_L(L-L_0)\frac{z}{L} - c_L\dot{L}\frac{z}{L}\right]\right)$$

Omitting Coriolis terms and adopting the notation $q = [x, y, z]^T$, with vectors and matrices in boldface, we can write the 3D bipedal SLIP dynamics as:

$$M(\ddot{\boldsymbol{q}} + \boldsymbol{g}) = \sum_{j=Right,Left}\left(-\left[k_L(L-L_0) + c\dot{L}\right]\frac{\boldsymbol{q}}{L} - \left[k_s\theta + c_s\dot{\theta}\right]\frac{\boldsymbol{q}}{L_s^2} - \left[k_f\phi + c_f\dot{\phi}\right]\frac{\boldsymbol{q}}{L_f^2}\right)_j, \qquad 4$$

where the summation corresponds to the right and left legs. The left-most brackets contain mechanisms that impart forces radially along the leg: $k_L$ is the leg stiffness, $L$ is the instantaneous leg length, $L_0$ is the leg resting length, $c_L$ is the leg damping, $\dot{L}$ is the instantaneous leg velocity. The middle bracket contains mechanisms that impart forces transverse to the leg axis in the sagittal plane: $k_s$ and $c_s$ are the sagittal-plane rotary stiffness and damping, respectively. $L_s$ denotes the sagittal-plane leg projection. Analogously in the right-most brackets, $k_f$ and $c_f$ represent the frontal-plane rotary stiffness and damping, respectively, and $L_f$ denotes the frontal-plane leg projection.

Note that we add gravitational acceleration vector, $\boldsymbol{g}$, to both sides of equation (4) to remove it from the right-hand side of the formulation. We can rewrite the dynamics as a linear map from the nonlinear transformations of the SLIP states to linear accelerations: $\ddot{\boldsymbol{q}} = \boldsymbol{\Theta}(\boldsymbol{q}, \dot{\boldsymbol{q}})\boldsymbol{\Xi}$,

$$\ddot{\boldsymbol{q}} + \boldsymbol{g} = \left(\begin{bmatrix} -\frac{k_L}{M} & \frac{k_L L_0}{M} & -\frac{c_L}{M} & \frac{k_s}{M} & \frac{c_s}{M} & \frac{k_f}{M} & \frac{c_f}{M} \end{bmatrix} \begin{bmatrix} x & y & z \\ \frac{x}{L} & \frac{y}{L} & \frac{z}{L} \\ \frac{\dot{L}x}{L} & \frac{\dot{L}y}{L} & \frac{\dot{L}z}{L} \\ \frac{\theta y}{L^2} & \frac{\theta x}{L^2} & 0 \\ \frac{\dot{\theta}y}{L^2} & \frac{\dot{\theta}x}{L^2} & 0 \\ 0 & 0 & \frac{\phi y}{L^2} \\ 0 & 0 & \frac{\dot{\phi}y}{L^2} \end{bmatrix}_{Right,Lleft}\right)^{\mathrm{T}} = \boldsymbol{\Theta}(\boldsymbol{q}, \dot{\boldsymbol{q}})\boldsymbol{\Xi} \qquad 5$$

The matrix $\boldsymbol{\Theta}(\boldsymbol{q}, \dot{\boldsymbol{q}}) \in \mathbb{R}^{3\times 14}$, is the function library containing candidate mechanisms describing COM accelerations, with seven candidate mechanisms per leg. Template Signatures were defined by the parameter matrix $\boldsymbol{\Xi}$.

#### Akaike Information Criterion (AIC)

In the following description of the AIC, we consider a nonlinear system with dynamics described by equation 5, where the estimated system dynamics are described by $\hat{f}(q)$, as the formulation below is not specific to models identified by Hybrid-SINDy.

The AIC is widely used to compare candidate models according to their number of free parameters (*i.e.*, model complexity), $k$, and log-likelihoods (*i.e.*, representativeness of the observations), $L(q, \hat{\mu})$. The AIC has the form:

$$AIC = 2k - 2\ln\big(L(q, \hat{\mu})\big). \qquad 6$$

Mangan and colleagues (2019) assumed that model errors are independently, identically, and normally distributed, enabling the AIC to be written in terms of the number of samples, $\rho$, and the sum of squared residuals [3]:

$$AIC = 2\tilde{k} + \rho \ln\left(\frac{\sum_{p=1}^{P}\sum_{i=1}^{I}\big(\hat{f}(\mathrm{q}) - \dot{q}\big)_i^{\,2}}{\rho}\right), \qquad 7$$

where in the outer summation, $P$ denotes the number of samples and in the inner summation, $I$ denotes the number of output states. A correction may be applied to the AIC formulation to account for bias due to finite sample sizes (AICc):

$$AICc = AIC + \frac{2(k+1)(k+2)}{P-k-2}. \qquad 8$$

The AICc score provides a level of support for each candidate model, with lower AICc scores indicating stronger support for a model. To compare competing models, a relative AICc score ($\Delta AICc$) may be calculated as the difference between the AICc score of the minimum-AICc model and that of competing models:

$$\Delta AICc_j = AICc_{min} - AICc_j, \qquad 9$$

where the subscript $j$, denotes the $j^{th}$ model. Burnham and Anderson [7] provided recommendations that $\Delta AICc \leq 2$ indicates strong support, while $\Delta AICc \leq 7$ indicates moderate support, $\Delta AICc$ values greater than seven have low support compared to competing models. Like Mangan and colleagues, we considered models with $\Delta AICc \leq 3$ to represent supported template signatures of walking [3].

### Comparison to alternative modeling frameworks

Here we present a brief analysis comparing Hybrid-SINDy to alternative modeling dimensionality reduction frameworks: Stepwise regression[8] and principal components analysis (PCA).

#### Stepwise regression

Stepwise regression is similar to Hybrid-SINDy in that it identifies sparse system models, often by eliminating or adding terms until terms no longer improve accuracy or are statistically significant. However, unlike Hybrid-SINDy, stepwise regression only selects a single model, even if multiple models are equally plausible representations of the system[9]. Further, stepwise regression is known to select different models depending on the data, especially when predictor variables are colinear[9,10]. However, it is unclear if stepwise regression could produce similar models as Hybrid-SINDy.

**Approach**: To test this, we replaced the SINDy algorithm with a similar approach using stepwise regression. Our stepwise regression approach is similar to that of Shamaei and Colleagues (2013), who estimated ankle quasi-stiffness from mechanics-based functional forms[8].

First, to provide an example of how stepwise regression produces different and less-sparse results than SINDy, we compared stepwise regression, as described in[8], to SINDy for the synthetic SLIP without noise and with moderate noise ($\eta$ = 1 mm). We report results for a single cluster in single-limb support.

Next, we applied a similar framework to human walking data and produced template signatures for walking in all three exoskeleton conditions, following the methodology described in the main manuscript.

**Results**: For synthetic SLIP data, Even in the noise-free condition, stepwise regression selected artificially high-dimensional dynamics (Table S1.1). The stepwise regression model had rank = 13, compared to the SINDy-based model, which identified 4 models ranging from 5- to 0-dimensional. Note that the SLIP model has at most 6 non-zero terms in each gait phase (*see Supplemental – S2*, *Table S2.1 for synthetic SLIP parameters*). With noise (Table S1.2), SINDy identified higher-dimensional solutions but also identified additional low-dimensional solutions that would likely be more plausible according to the AIC.

Table S1.1: Noise-free synthetic SLIP template signatures in one cluster in single-limb support.

| Coefficient | Stepwise solution (rank = 13) | SINDy – model 1 (rank = 5) | SINDy – model 2 (rank = 4) | SINDy – model 3 (rank = 2) | SINDy – model 4 (rank = 0) |
|---|---|---|---|---|---|
| $k_{L,R}$ | -244.8 | -260.8 | -221.7 | -385.6 | 0 |
| $k_{L,L}$ | -359.4 | -359.6 | -345.1 | 0 | 0 |
| $k_{L,R}L_{0,R}$ | 244.0 | 257.6 | 224.8 | 361.3 | 0 |
| $k_{L,R}L_{0,L}$ | 252.5 | 251.9 | 223.1 | 0 | 0 |
| $c_{L,R}$ | -0.9 | 0 | 0 | 0 | 0 |
| $c_{L,L}$ | -14.9 | -15.1 | 0 | 0 | 0 |
| $k_{s,R}$ | 0.8 | 0 | 0 | 0 | 0 |
| $k_{s,L}$ | -0.6 | 0 | 0 | 0 | 0 |
| $c_{s,R}$ | 0.0 | 0 | 0 | 0 | 0 |
| $c_{s,R}$ | 0 | 0 | 0 | 0 | 0 |
| $k_{f,R}$ | 0.8 | 0 | 0 | 0 | 0 |
| $k_{f,L}$ | -0.6 | 0 | 0 | 0 | 0 |

| | | | | | |
|---|---|---|---|---|---|
| $c_{f,R}$ | -0.0 | 0 | 0 | 0 | 0 |
| $c_{f,R}$ | 0.0 | 0 | 0 | 0 | 0 |

Table S1.2: Synthetic SLIP template signatures with noise ($\eta = 1mm$) in one cluster in single-limb support.

| Coefficient | Stepwise solution (rank = 11) | SINDy – model 1 (rank = 12) | SINDy – model 2 (rank = 11) | SINDy – model 3 (rank = 8) | SINDy – model 4 (rank = 3) |
|---|---|---|---|---|---|
| $k_{L,R}$ | -455.0 | -407.8 | -219.4 | 32.6 | 41.0 |
| $k_{L,L}$ | 0 | -31.9 | -112.7 | -299.9 | -269.0 |
| $k_{L,R}L_{0,R}$ | 797.6 | 729.4 | 459.2 | 18.6 | 0 |
| $k_{L,R}L_{0,L}$ | -373.6 | -325.3 | -171.8 | 179.2 | 170.8 |
| $c_{L,R}$ | 0 | 0 | 0 | 0 | 0 |
| $c_{L,L}$ | -8.8 | -8.0 | 0 | 0 | 0 |
| $k_{s,R}$ | -306.8 | -283.5 | -192.7 | 0 | 0 |
| $k_{s,L}$ | 217.5 | 199.4 | 128.4 | 0 | 0 |
| $c_{s,R}$ | -4.2 | -4.2 | -4.1 | -3.8 | 0 |
| $c_{s,R}$ | -9.2 | -9.3 | -9.3 | -8.8 | 0 |
| $k_{f,R}$ | -338.9 | -315.6 | -224.3 | -51.9 | 0 |
| $k_{f,L}$ | 260.6 | 242.2 | 169.8 | 0 | 0 |
| $c_{f,R}$ | -17.3 | -17.4 | -17.9 | -15.3 | 0 |
| $c_{f,R}$ | 0 | 0 | 0 | 0 | 0 |

In human data, stepwise regression also identified rank = 13 dynamics, while SINDy identified lower-rank dynamics (Table S1.3). Consistent with single-cluster results in the synthetic SLIP, stepwise regression-based template signatures were not sparse (Figure S1.2; compare to Figure 3 in the main manuscript). Figure S1.3 shows that the post-stroke template signatures are also not sparse, making it unclear which mechanisms are most important for template representations of CoM dynamics (compare to Figure 6 in the main manuscript).

Table S1.3: Example human template signatures in one cluster in single-limb support.

| Coefficient | Stepwise solution (rank = 13) | SINDy – model 1 (rank = 12) | SINDy – model 2 (rank = 6) | SINDy – model 3 (rank = 2) | SINDy – model 4 (rank = 0) |
|---|---|---|---|---|---|
| $k_{L,R}$ | -79.1 | -65.3 | -99.2 | -150.6 | 0 |
| $k_{L,L}$ | 18.9 | 5.1 | 11.8 | 0 | 0 |
| $k_{L,R}L_{0,R}$ | 43.5 | 15.7 | 75.8 | 132.0 | 0 |
| $k_{L,R}L_{0,L}$ | 10.1 | 35.1 | 0 | 0 | 0 |
| $c_{L,R}$ | 0 | 0 | 0 | 0 | 0 |
| $c_{L,L}$ | -1.4 | 0 | 0 | 0 | 0 |
| $k_{s,R}$ | 22.4 | 34.1 | 0 | 0 | 0 |
| $k_{s,L}$ | -19.3 | -31.9 | -13.8 | 0 | 0 |
| $c_{s,R}$ | -1.3 | -0.7 | 1.1 | 0 | 0 |
| $c_{s,R}$ | -0.5 | -0.3 | 0 | 0 | 0 |

| | | | | | |
|---|---|---|---|---|---|
| $k_{f,R}$ | 29.6 | 44.4 | 16.5 | 0 | 0 |
| $k_{f,L}$ | -20.2 | -31.3 | -9.2 | 0 | 0 |
| $c_{f,R}$ | -1.0 | 0 | 0 | 0 | 0 |
| $c_{f,R}$ | 0 | 0 | 0 | 0 | 0 |

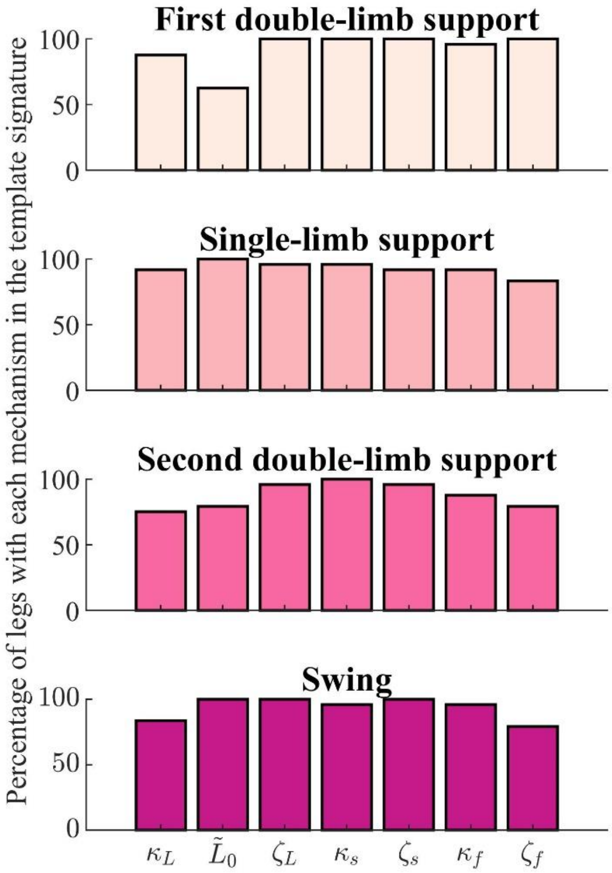


**Figure S1.2: The percentage of unimpaired legs whose stepwise regression-based template signatures contained each mechanism in each gait phase.** For 24 legs, the percentage of legs for which each template signature mechanism was selected by the Hybrid-SINDy algorithm. Colors denote each gait phase. Mechanisms selected in a larger percentage of legs suggest common representations of CoM dynamics, while less frequently selected mechanisms reflect individual-specific template features describing CoM dynamics.

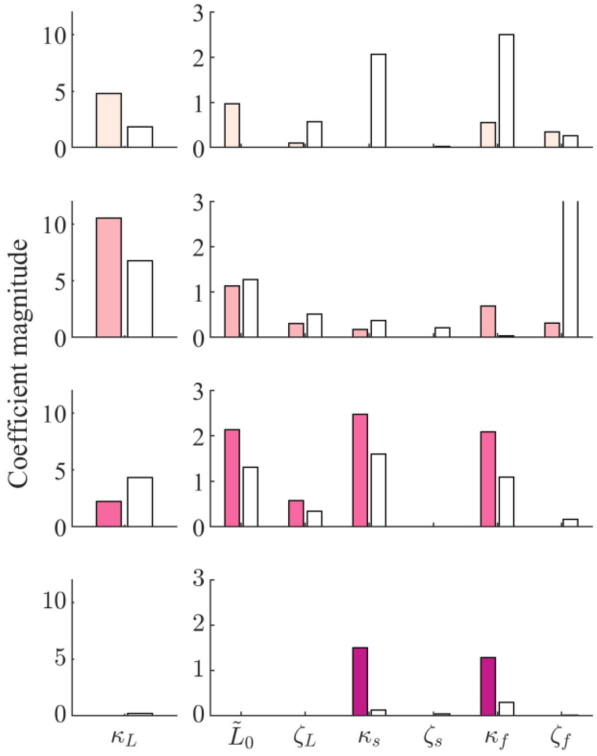


**Figure S1.3: Inter-leg differences in template signatures in one stroke survivor identified using stepwise regression.** Non-paretic (colored bars) and paretic (white bars) template signatures for one individual with post-stroke hemiparesis. Each plot represents a different gait phase.

**Interpretation**:
These results showcase that stepwise regression tends to select less parsimonious models than Hybrid-SINDy, hindering interpretation of template signatures. While regression hyperparameters may be tuned to reduce this dimensionality, Hybrid-SINDy's use of clustering and model selection within clusters likely enables more sparse template representations of CoM dynamics to be identified. Therefore, Hybrid-SINDy was appropriate for our work.

#### Principal components analysis (PCA)

Principal component analysis (PCA) could identify interpretable basis vectors describing CoM dynamics. For example, we could apply (e.g.) PCA or NMF to the nonlinear template states and use information criteria to select an optimal number of modes in the model[11].

However, PCA would not satisfy our goal of interpreting individual differences in template-based representations of CoM dynamics. PCA-based models would not be sparse in the space of template parameters: If the optimal (*e.g.*, according to the AIC) PCA or NMF-based model was unidimensional, the first PC would contain multiple template states. The higher-dimensional modes of these models would likely also contain multiple non-zero template states. Therefore, PCA would not provide the same mechanically intuitive interpretability as the sparse regression of template states used by Hybrid-SINDy.

**Approach**: To exemplify the limitation of PCA (with potential generalization to other dimensionality reduction approaches), we show that a sparse PCA-based model is still dense in the template mechanism space. Using data from our synthetic SLIP, we applied PCA to the template states, then used SINDy to identify the linear map, $\boldsymbol{\Xi}_{\mathbf{PC}}$, from PCs to CoM accelerations. We then projected the map back into the original template basis ($\boldsymbol{\Xi}_{\boldsymbol{template}}$ in Table S1.4**).**

**Results**: Table S1.4 shows an example of a 1-D linear map identified by SINDy. While the linear map is sparse in the PC space, it becomes dense when projected back into the original template mechanism space (Table S1.4).

Table S1.4: 1-dimensional PCA-based model of CoM dynamics.

| **PCA model term** | **1-D PC model ($\boldsymbol{\Xi}_{\mathbf{PC}}$)** | **1-D PC model projected into template state space ($\boldsymbol{\Xi}_{\boldsymbol{template}}$)** |
|---|---|---|
| 1 | 0 | -0.417 |
| 2 | 0 | -0.323 |
| 3 | -0.351 | -0.418 |
| 4 | 0 | -0.323 |
| 5 | 0 | -0.393 |
| 6 | 0 | 0.323 |
| 7 | 0 | -0.136 |
| 8 | 0 | 0.172 |
| 9 | 0 | -0.133 |
| 10 | 0 | 0.125 |
| 11 | 0 | -0.185 |
| 12 | 0 | 0.106 |
| 13 | 0 | -0.142 |
| 14 | 0 | -0.183 |

**Interpretation**: Because—even for low-dimensional PC-based models—the resulting template model is full-dimensional, we cannot readily interpret which well-understood mechanisms are most important for describing CoM dynamics using template models. This limitation would likely generalize to other techniques like non-negative matrix factorization[12].
