## Supplemental - S2 for "Quantifying changes in individual-specific template-based representations of center-of-mass dynamics during walking with ankle exoskeletons using Hybrid-SINDy"

#### Overview

This supplemental provides details of the simulations using a synthetic spring-loaded inverted pendulum (SLIP) dynamics to validate the Hybrid Sparse Identification of Nonlinear Dynamics (Hybrid-SINDy) for human walking.

1. Validating Hybrid-SINDy using a synthetic SLIP walking model
    a. Effects of measurement noise on algorithm performance
    b. Effects of missing physics on algorithm performance
2. Evaluation of collinearity of template signature state variables

### Validating Hybrid-SINDy using a synthetic SLIP walking model

We evaluated Hybrid-SINDy's ability to accurately identify and select walking dynamics, using synthetic data generated from simulations of a bipedal SLIP walker with known dynamics (Table S2.1 and Figure S2.1)[1]. The bipedal SLIP had dynamics matching those used in human data (*Supplemental – S1*), with the addition of foot mass to enable simulation of a full gait cycle. The synthetic SLIP had asymmetric leg stiffness and damping, and masses representing the pelvis ($M$ = 56 kg) and feet ($M_f$ = 7 kg) (Table S2.1; Figure S2.1). We omitted rotary mechanisms from the SLIP dynamics but used the full function library described in *Supplemental – S1*, equation 4. To replicate the human walking datasets, we simulated 120 gait cycles each from randomly perturbed initial conditions with an average initial velocity of 1.20 m/s. The SLIP walked at approximately 1 stride/s.

Table S2.1 – Bipedal SLIP normalized simulation parameters.

| Term | ***[Right leg, Left leg]*** | | | |
|---|---|---|---|---|
| | **$DS_1$** | **SS** | **$DS_2$** | **SW** |
| $\kappa_L$ | [10.8, 9.1] | [25.1, 18.82] | [9.03, 10.68] | [36.6,25.3] |
| $\tilde{L}_0$ | [1.0, 1.0] | [1.0, 0.7] | [1.0, 1.0] | [0.7, 0.7] |
| $\zeta_L$ | [0.04, -0.07] | [-0.03, -0.04] | [0, 0] | [-0.40, -0.57] |
| $\kappa_s$ | 0 | 0 | 0 | 0 |
| $\zeta_s$ | 0 | 0 | 0 | 0 |
| $\kappa_f$ | 0 | 0 | 0 | 0 |
| $\zeta_f$ | 0 | 0 | 0 | 0 |

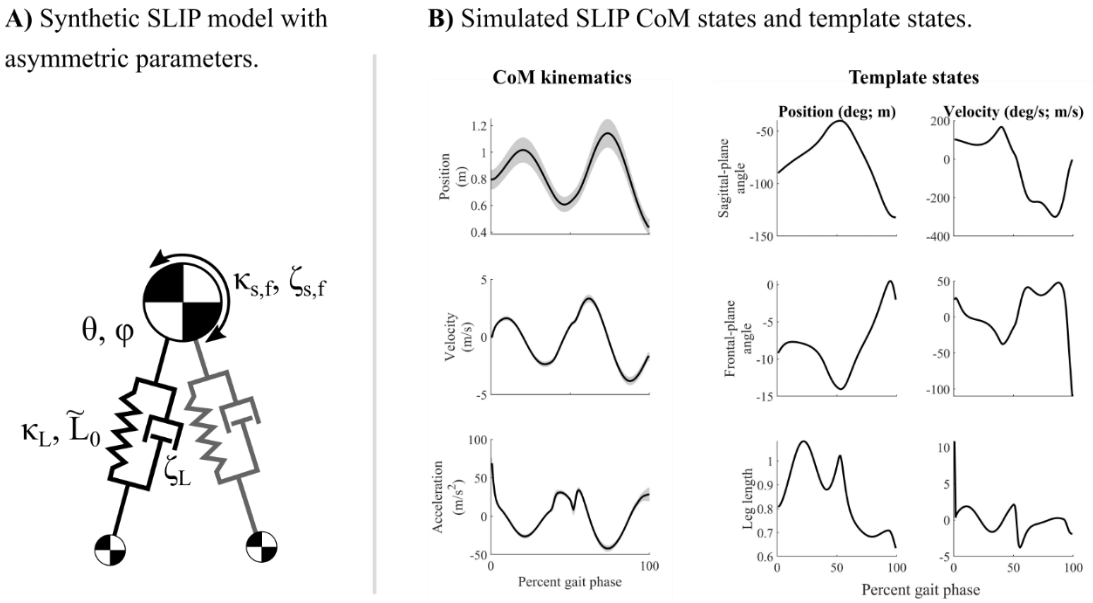


**Figure S2.1: Synthetic SLIP model depiction and simulated CoM states.** A) Two-dimensional depictions of the simulated SLIP. The simulated SLIP used leg springs and dampers, as well as foot masses to simulate full strides. B) Time-series measurements of simulated CoM position, velocity, and acceleration for the simulated SLIP (left; average ± 1SD). Template states (right) show trajectories of the template variables used in the function library to estimate template signature coefficients (average ± 1SD).

We conducted two analyses using the simulations of a synthetic SLIP model: First, we evaluated the effects of measurement noise on Hybrid-SINDy's ability to estimate synthetic SLIP parameters in the presence of sensor noise. Second, we evaluated Hybrid-SINDy's ability to estimate synthetic SLIP parameters when a subset of the terms constituting the true system dynamics is missing from the function library.

#### Effects of measurement noise on algorithm performance

**Approach**: As SINDy's performance is sensitive to sensor noise, we identified template signatures using simulation measurements with Gaussian noise ranging logarithmically from 10 *nm* to 10 *cm* added to the position measurements[2]. After adding noise, we differentiated each gait cycle separately before concatenating the strides to avoid errors in the estimated time derivatives due to changes in the initial conditions of each simulation. We quantified the Hybrid-SINDy's ability to reconstruct model dynamics using the number of correctly selected mechanisms and the accuracy of estimated coefficients. We also evaluated the ability of each model to predict the ground truth (noise-free) CoM accelerations.

**Results**: In the noise-free condition, the Hybrid-SINDy algorithm correctly identified the salient features of the simulated SLIP's template signature during single-limb support (**Figure S2.2**). Specifically, Hybrid-SINDy identified non-zero leg stiffness ($\kappa_L$), resting length ($\tilde{L}$), and damping ($\zeta_L$) terms only – consistent with the SLIP's dynamics. In the noise-free condition, 86% of terms were correctly selected and only one term, leg damping in first double-limb support, was incorrectly selected (**Figure S2.2**B). Both double-limb support phases were very short in the simulated SLIP: 1.3% and 4.6% of samples in first and second double-limb support – smaller than the 7.4% of the training data in each cluster. Consequently, all clusters with centroids in the double-limb support phases contained samples spanning double- and single-limb support. Hybrid-SINDy incorrectly omitted only damping terms with small coefficients (**Figure S2.2**). In single-limb support and swing, coefficients were within 2% of the ground truth values. While prediction accuracy (averaged across output dimensions) was lower than expected ($r^2 = 0.88$), errors occurred primarily at the transitions between gait phases and in the double-limb support phases (**Figure S2.2**A, left: deviations in the predictions (colors) relative to ground truth simulation results (gray); transitions occur where colors change).

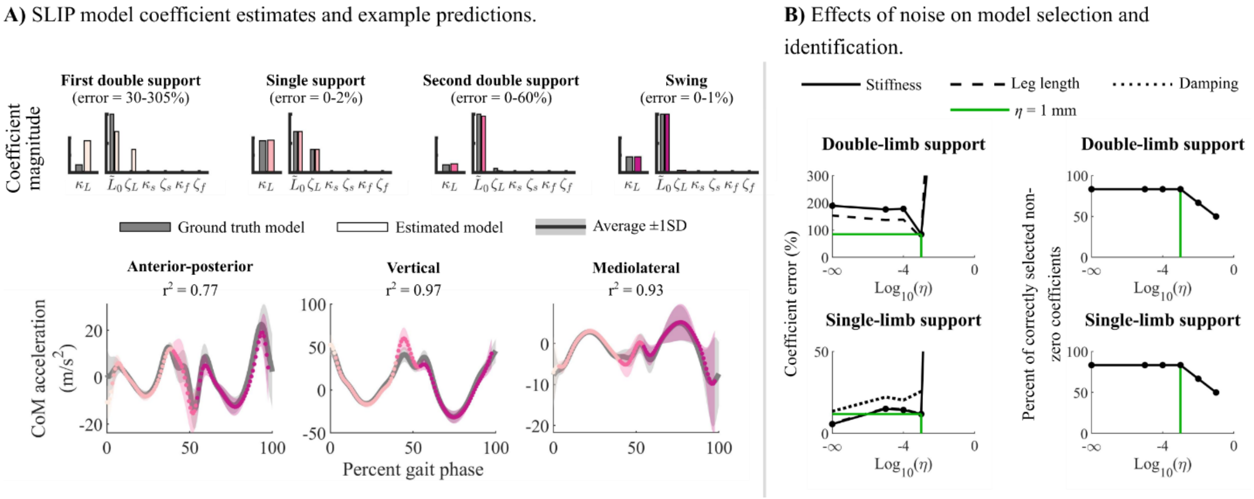


**Figure S2.2: Parameter estimates, prediction accuracy of simulated CoM kinematics, and the effects of noise on model identification accuracy.** A) Model coefficients (top) can be compared between walking conditions. Bar plots show ground truth synthetic SLIP parameters (gray) and the corresponding estimated template signature coefficients (colors). The range of percent errors in coefficients is shown above each plot. The average (±1SD) ground truth CoM accelerations of the synthetic SLIP (bottom; gray) and model predictions in each gait phase (colors). Model accuracies are quantified using the coefficient of determination ($r^2$). B) Effects of noise on model coefficient percent error (left) and the percent of correctly selected mechanisms (right). All horizontal axes represent noise magnitudes on a log scale. For coefficient error, percent errors were computed for stiffness (solid line), leg length (dashed line), and damping (dotted line) separately. Green lines denote the $\eta$ = 1 mm noise level.

When measurement noise with a standard deviation of 1 mm—similar to that of our motion capture system—was added to the system measurements, Hybrid-SINDy correctly selected 83% of mechanisms, with incorrect mechanisms selection occurring primarily in double-limb support. Template signature coefficients were estimated with an average error of less than 29% in single-limb support and swing (**Figure S2.2**). In single-limb support, Hybrid-SINDy was still able to estimate large coefficients (*i.e.*, leg stiffness and resting length) with less than 1% error. Even at this noise level, the estimated template signature could still predict the noise-free data with similar accuracy to the noise-free data ($r^2$ = 0.83). The accuracy of coefficient estimates and model predictions rapidly declined at noise levels with standard deviations larger than 1 mm.

**Implications**: Hybrid-SINDy's ability to identify template signatures from forward simulations of a synthetic SLIP model with known dynamics at measurement noise levels comparable to marker-based motion capture suggests that individual differences in template signatures of human gait data likely reflect some differences in the underlying CoM dynamics. Hybrid-SINDy performed as expected in the noise-free condition, accurately identifying single-limb support and swing dynamics, except for small damping terms. Terms with small contributions to system behavior, such as damping in this study, may be omitted by the AICc in favor of parsimony[3,4]. Therefore, Hybrid-SINDy will not always select the true system dynamics but will select terms critical to describing the salient aspects of CoM dynamics. In our human walking data, Hybrid-SINDy may, therefore, omit mechanisms describing subtle idiosyncrasies in CoM dynamics that could be important for gait stability or efficiency. Future experiments using different constraints or perturbations may identify additional mechanisms needed to describe CoM dynamics.

The reduced accuracy of synthetic SLIP template signatures in double-limb support and with increasing noise magnitude is consistent with the results of Hybrid-SINDy applied to a spring-mass hopper[5]. However, accurate estimates of single-limb support CoM dynamics and longer double-limb support times in human walking data support our use of 800 samples per cluster for human template signatures. Longer time-series datasets would help to ensure that clusters are small enough to span only a single gait phase but large enough to remain robust to sensor noise.

#### Effects of missing physics on algorithm performance

**Approach**: We tested the robustness of estimated template signature coefficients using simulations of a synthetic SLIP: We simulated SLIP walking at speeds ranging from 1.00-1.75 m/s (Froude speeds: 0.10-0.31) using identical dynamics across all simulations. Next, we confirmed that, in the noise-free condition, Hybrid-SINDy identified the same model coefficients for all simulations when using the *complete* function library that contained all functional forms constituting the SLIP's dynamics. To characterize how missing functional forms affected Hybrid-SINDy's ability to identify model coefficients, we applied Hybrid-SINDy to the same data but with the left- and right-leg axial damping terms removed from the function library. We quantified the effect of missing physics (damping terms) as the percent differences in the estimated model coefficients (averaged across coefficients) between simulations. We defined the similarity of kinematics between simulations as the root-mean-squared (RMS) difference between phase-averaged kinematics.

**Results:** The kinematic differences observed in SLIP simulations were larger than those seen in our human data (Figure S2.3A, shaded boxes represent the range of human data). We observed this difference despite participants walking at self-selected speeds ranging from 1.20-1.50 m/s (Froude speeds: 0.16-0.29), comparable to the range tested *in silico*. For *within-participant* comparisons, differences in kinematics were smaller than those observed in any simulation pairs (Figure S2.3A, blue shaded region; Figure S2.3C).

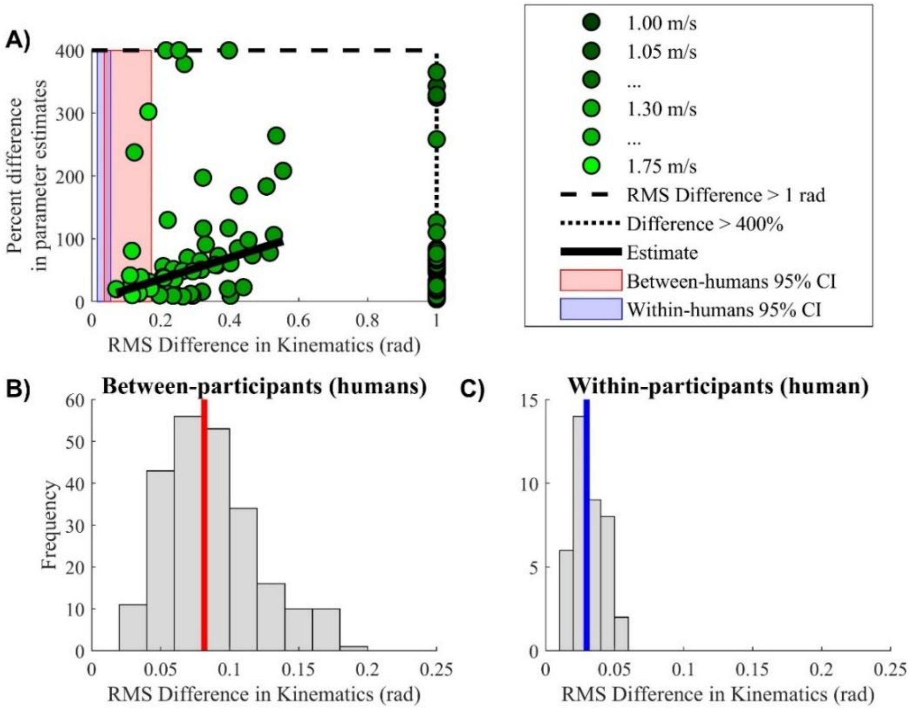


**Figure S2.3: Regression analysis of differences in estimated synthetic SLIP coefficients as a function of differences in kinematics, and compared to differences in human kinematics**. Pair-wise differences in estimated coefficients and kinematics between simulations was computed as the root-mean-squared difference across coefficients or the stride-averaged kinematic trajectories, respectively. Note that the model coefficients were normalized, such that the differences in coefficients are dimensionless. Also note that we fixed template signature structure to match that of the signature selected for the 1.35 m/s simulation. A) Regression analysis of synthetic data using the 10 fastest speeds (1.30-1.75 m/s). Green dots denote simulation pairs. Some simulation pairs had large differences due to near-zero coefficient estimates; their values are truncated on the vertical axis at 400% (dashed line). Simulations at speeds under 1.30 m/s had large differences in kinematics; their values are truncated on the horizontal axis (dotted line). The shaded regions represent 95% confidence intervals of differences in kinematics in the human data between- (red) and within-participants (blue). These colors correspond to the colors of vertical lines in panels B and C. B) Histogram of between-participants differences in kinematics in the human data. Each sample represents a pair of participants in the same walking condition. The red line represents the median difference in kinematics. C) Histogram of within-participants differences in kinematics in human data. Each sample represents a pair of exoskeleton conditions (e.g., shoes-only vs. zero-stiffness) for the same participant. The blue line represents the median difference in kinematics.

When the *complete* function library was included, Hybrid-SINDy identified nearly consistent template signatures across all walking speeds. In single-limb support and swing, the estimated coefficients were consistent across simulations; differences in leg stiffness and resting length estimates were 3.9 ± 4.5% and 0.7 ± 0.9%. Similar to our analysis of noise, estimated coefficients were more variable in double-limb support (differences > 269%), likely because the clusters were larger than the simulated double-limb support phases.

Compared to using the complete function library to identify template signatures, *missing physics* resulted in larger differences in estimated coefficients across the full speed range. When the same term was active in two simulations, the differences in leg stiffness and resting length estimates were 6.3 ± 5.2% and 1.3 ± 1.4%, respectively, in single-limb support. Unsurprisingly, differences were larger in double-limb support (>>25%) as double-limb support phases were short compared to the cluster sizes. Different template signature structures (i.e., different non-zero terms) were selected across simulations, making this analysis similar to a *between-participants* comparison.

Because our analysis of human data used a single (shoes-only) template signature structure for *within-participants* comparisons, we conducted a second analysis to reflect this comparison: We fixed the synthetic SLIP template signature structures to the structure selected for an intermediate speed (1.35 m/s) and estimated model coefficients. Compared to template signatures with unique structures identified for each simulation, the differences in axial leg stiffness and resting length estimates were only marginally reduced: 5.7 ± 5.0% and 0.5 ± 0.5%, respectively, in single-limb support and remained larger in double-limb support.

In this within-participants analysis, missing physics resulted in differences in coefficients driven by changes in simulated kinematics. At speeds slower than 1.30 m/s, differences in the estimated coefficients were not correlated with differences in kinematics.  However, within the 1.30-1.75 m/s speed range, larger differences in simulated kinematics were associated with larger differences in coefficient estimates ($r^2 = 0.50$, $p < 0.001$; Figure S2.3A, black estimate line).

**Implications**: Small differences in kinematics in humans compared to the synthetic SLIP simulations suggest that large differences in template signature coefficients are unlikely due entirely to differences in kinematics. However, at least 5-10% differences in template signature coefficients can be attributed to changes in kinematics rather than CoM dynamics. Larger differences in coefficient estimates may be needed for human data, depending on how well the model captures the underlying dynamics.

### Covariation of template signature state variables

Covariation among the state variables corresponding to mechanisms in the Hybrid-SINDy function library can hinder the interpretation of template signatures. Specifically, covariation in state variables can alter coefficient estimates if both mechanisms corresponding to two co-varying states are selected, or if redundant mechanisms are selected at random for different participants.

**Approach**: To investigate the effect of covariation between state variables for the human data, we correlated SLIP states within each gait phase separately (**Figure S2.4**; average coefficient of determination across participants).

**Results**: Leg stiffness and resting length were most strongly correlated, as expected ($r^2 = 0.99$). These states were both selected by Hybrid-SINDy. Most correlations were less than $r^2 = 0.70$ (**Figure S2.5** highlights strong correlations = $r^2 > 0.70$). While frontal-plane rotary stiffness ($\kappa_f$) and damping ($\zeta_f$) in double-limb support were strongly correlated, only frontal-plane rotary stiffness was selected for a subset of participants (see Figure 3, main manuscript).

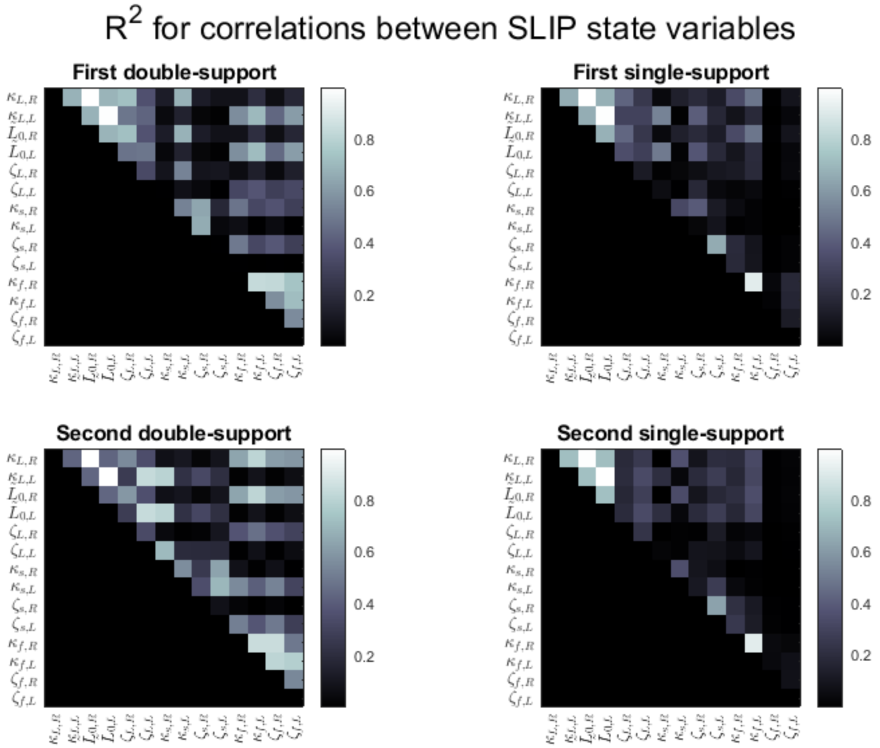


**Figure S2.4: Correlations between SLIP states in human walking data for each gait phase**. Colormaps of R-squared ($R^2$) values denoting correlations between each pair of state variables in each gait phase (one panel for each phase, denoted by titles above each panel). Black boxes denote $R^2 = 0$ and white boxes denote $R^2 = 1$. The lower triangle is omitted for clarity (the correlation matrix is symmetric). Along the axes, the second subscript on each variable denotes the right (R) or left (L) leg. Only axial stiffness ($\kappa_L$) and leg resting length ($L_0$) states were, as expected, very strongly correlated.

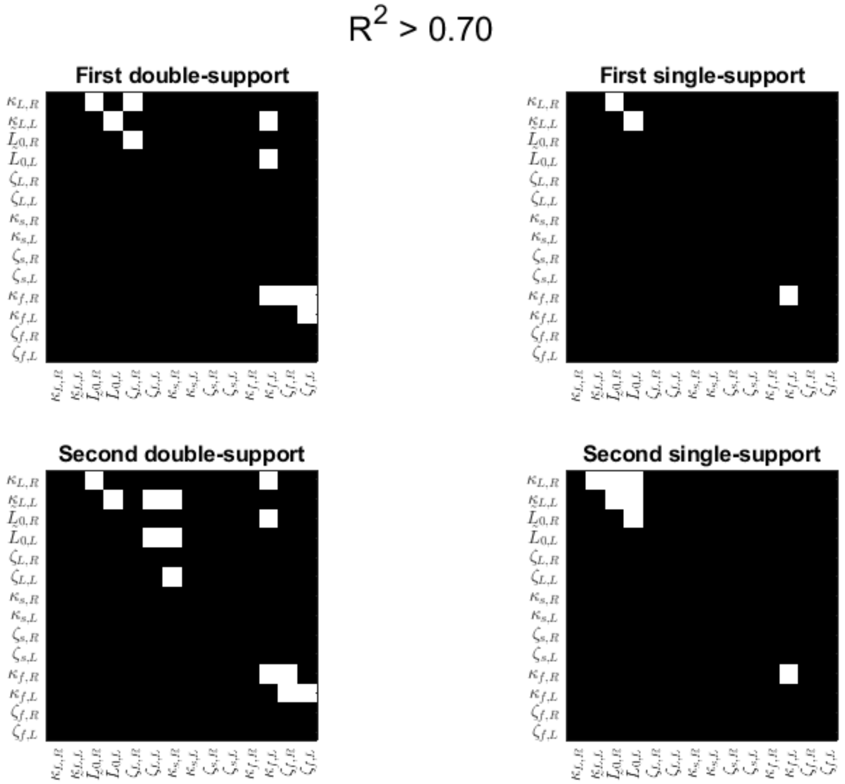


**Figure S2.5: Strong correlations between SLIP states in human walking data for each gait phase**. Colormaps of R-squared ($R^2$) values highlight strong correlations ($R^2 > 0.70$) between each pair of state variables in each gait phase (one panel for each phase, denoted by titles above each panel). Black boxes denote $R^2 = 0$ and white boxes denote $R^2 > 0.70$. The lower triangle is omitted for clarity (the correlation matrix is symmetric). Along the axes, the second subscript on each variable denotes the right (R) or left (L) leg. While frontal-plane rotary stiffness ($\kappa_f$) and damping ($\zeta_f$) in double-limb support were strongly correlated, only frontal-plane rotary stiffness was selected for some participants. However, Hybrid-SINDy consistently selected frontal-plane rotary stiffness, suggesting that stronger correlations are needed to select different mechanisms for different participants.

**Interpretation**: Hybrid-SINDy's performance appears robust to moderate correlations in the state variables. Covariation in the state variables did not result in unexpected variables being included in the template signatures. However, different combinations of parameters could have similar statistical plausibility. In this case, our use of the Akaike Information Criterion and multi-model inference acknowledges the existence of multiple plausible template signatures[3,4]. While we selected mechanisms with distinct relationships to CoM accelerations, studies using larger or non-mechanistic function libraries may be more strongly impacted by covariation in the state variables. Such studies should evaluate covariation among state variables before model fitting.
